# Ancient origin of an urban underground mosquito

**DOI:** 10.1101/2025.01.26.634793

**Authors:** Yuki Haba, Matthew L. Aardema, Maria O. Afonso, Natasha M. Agramonte, John Albright, Ana Margarida Alho, Antonio P.G. Almeida, Haoues Alout, Bulent Alten, Mine Altinli, Raouf Amara Korba, Stefanos S. Andreadis, Vincent Anghel, Soukaina Arich, Arielle Arsenault-Benoit, Célestine Atyame, Fabien Aubry, Frank W. Avila, Diego Ayala, Rasha S. Azrag, Lilit Babayan, Allon Bear, Norbert Becker, Anna G. Bega, Sophia Bejarano, Ira Ben-Avi, Joshua B. Benoit, Saïd C. Boubidi, William E. Bradshaw, Daniel Bravo-Barriga, Rubén Bueno-Marí, Nataša Bušić, Viktoria Čabanová, Brittany Cabeje, Beniamino Caputo, Maria V. Cardo, Simon Carpenter, Elena Carreton, Mouhamadou S. Chouaïbou, Michelle Christian, Maureen Coetzee, William R. Conner, Anton Cornel, C. Lorna Culverwell, Aleksandra I. Cupina, Katrien De Wolf, Isra Deblauwe, Brittany Deegan, Sarah Delacour-Estrella, Alessandra della Torre, Debora Diaz, Serena E. Dool, Vitor L dos Anjos, Sisay Dugassa, Babak Ebrahimi, Samar Y.M. Eisa, Nohal Elissa, Sahar A.B. Fallatah, Ary Faraji, Marina V. Fedorova, Emily Ferrill, Dina M. Fonseca, Kimberly A. Foss, Cipriano Foxi, Caio M. França, Stephen R. Fricker, Megan L. Fritz, Eva Frontera, Hans-Peter Fuehrer, Kyoko Futami, Enas H.S. Ghallab, Romain Girod, Mikhail I. Gordeev, David Greer, Martin Gschwind, Milehna M. Guarido, Teoh Guat Ney, Filiz Gunay, Eran Haklay, Alwia A.E. Hamad, Jun Hang, Christopher M. Hardy, Jacob W. Hartle, Jenny C. Hesson, Yukiko Higa, Christina M. Holzapfel, Ann-Christin Honnen, Angela M. Ionica, Laura Jones, Përparim Kadriaj, Hany A. Kamal, Colince Kamdem, Dmitry A. Karagodin, Shinji Kasai, Mihaela Kavran, Emad I.M. Khater, Frederik Kiene, Heung-Chul Kim, Ilias Kioulos, Annette Klein, Marko Klemenčić, Ana Klobučar, Erin Knutson, Constantianus J.M. Koenraadt, Linda Kothera, Pauline Kreienbühl, Pierrick Labbé, Itay Lachmi, Louis Lambrechts, Nediljko Landeka, Christopher H. Lee, Bryan D. Lessard, Ignacio Leycegui, Jan O. Lundström, Yoav Lustigman, Caitlin MacIntyre, Andrew J. Mackay, Krisztian Magori, Carla Maia, Colin A. Malcolm, Ralph-Joncyn O. Marquez, Dino Martins, Reem A. Masri, Gillian McDivitt, Rebekah J. McMinn, Johana Medina, Karen S. Mellor, Jason Mendoza, Enrih Merdić, Stacey Mesler, Camille Mestre, Homer Miranda, Martina Miterpáková, Fabrizio Montarsi, Anton V. Moskaev, Tong Mu, Tim W.R. Möhlmann, Alice Namias, Ivy Ng’iru, Marc F. Ngangué, Maria T. Novo, Laor Orshan, José A. Oteo, Yasushi Otsuka, Rossella Panarese, Claudia Paredes-Esquivel, Lusine Paronyan, Steven T. Peper, Dušan V. Petrić, Kervin Pilapil, Cristina Pou-Barreto, Sebastien J. Puechmaille, Ute Radespiel, Nil Rahola, Vivek K Raman, Hamadouche Redouane, Michael H. Reiskind, Nadja M. Reissen, Benjamin L. Rice, Vincent Robert, Ignacio Ruiz-Arrondo, Ryan Salamat, Amy Salamone, M’hammed Sarih, Giuseppe Satta, Kyoko Sawabe, Francis Schaffner, Karen E. Schultz, Elena V. Shaikevich, Igor V. Sharakhov, Maria V. Sharakhova, Nader Shatara, Anuarbek K. Sibataev, Mathieu Sicard, Evan Smith, Ryan C. Smith, Nathalie Smitz, Nicolas Soriano, Christos G. Spanoudis, Christopher M. Stone, Liora Studentsky, Tatiana Sulesco, Luciano M. Tantely, La K. Thao, Noor Tietze, Ryan E. Tokarz, Kun-Hsien Tsai, Yoshio Tsuda, Nataša Turić, Melissa R. Uhran, Isik Unlu, Wim Van Bortel, Haykuhi Vardanyan, Laura Vavassori, Enkelejda Velo, Marietjie Venter, Goran Vignjević, Chantal B.F. Vogels, Tatsiana Volkava, John Vontas, Heather M. Ward, Nazni Wasi Ahmad, Mylène Weill, Jennifer D. West, Sarah S. Wheeler, Gregory S. White, Nadja C. Wipf, Tai-Ping Wu, Kai-Di Yu, Elke Zimmermann, Carina Zittra, Petra Korlević, Erica McAlister, Mara K.N. Lawniczak, Molly Schumer, Noah H. Rose, Carolyn S. McBride

## Abstract

Understanding how life is adapting to urban environments represents an important challenge in evolutionary biology. Here we investigate a widely cited example of urban adaptation, *Culex pipiens* form *molestus*, also known as the London Underground Mosquito. Population genomic analysis of ∼350 contemporary and historical samples counter the popular hypothesis that *molestus* originated belowground in London less than 200 years ago. Instead, we show that *molestus* first adapted to human environments aboveground in the Middle East over the course of >1000 years, likely in concert with the rise of agricultural civilizations. Our results highlight the role of early human society in priming taxa for contemporary urban evolution and have important implications for understanding arbovirus transmission.

## Main

The rise of modern cities is rapidly reshaping our planet and imposing novel selective pressure on the living organisms around us. Many species have begun to adapt to these unique challenges. A review of the literature highlights at least 130 examples of animals, plants, and microbes that have evolved responses to dense urban environments (*1*). Yet how such adaptations occur and the amount of time they require remain poorly understood. As urbanization accelerates over the coming decades (*2*), there is a pressing need to better understand the mechanisms and timescale of urban adaptation.

One of the most widely cited examples of urban adaptation involves the northern house mosquito *Culex pipiens* Linnaeus, 1758 (Fig. 1A). *Cx. pipiens* is common in temperate zones across the world (*3*, *4*). In Europe and North America, an ancestral form has long been appreciated as a bird-biting mosquito that requires open space for mating and pauses reproduction (i.e. diapauses) during the cold northern winter (Fig. 1B–C, blue) (*3*). However, a derived, human-biting form thrives in urban belowground environments, such as subways, cellars, and cesspits, and differs from its aboveground counterpart in ways that seem perfectly suited to subterranean life (Fig. 1B–C, red) (*3*). The belowground mosquitoes are able to mate in confined spaces and remain active in winter. Adult females readily bite humans (and other mammals). Yet if hosts are scarce, they can develop a first clutch of eggs without taking any blood, a trait known as autogeny. Strikingly, despite this array of genetically based behavioral and physiological differences, the two mosquitoes show no consistent morphological differences (*3*). They are formally considered distinct forms: the bird-biting *Cx. pipiens* f. *pipiens* Linnaeus, 1758 and the human-biting *Cx. pipiens* f. *molestus* Forskål, 1775 (*5*), hereafter referred to as simply *pipiens* and *molestus*, respectively.

**Figure 1.**
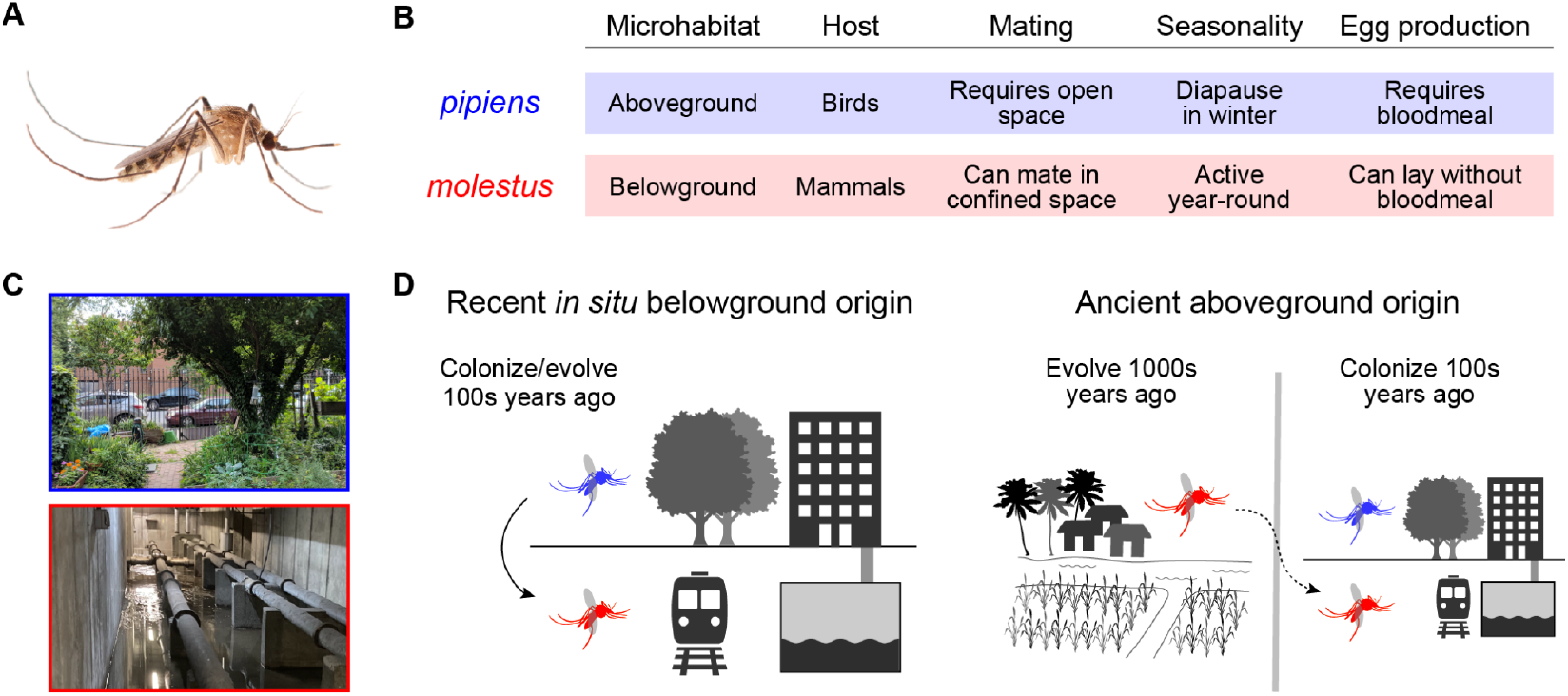
*Culex pipiens* form *molestus* behavior, ecology, and hypothetical origin. (**A**) Female *Cx. pipiens* complex mosquito. (**B**) Behavioral and physiological characteristics of *Cx. pipiens* forms in northern Eurasia. At warmer latitudes *molestus* can breed aboveground. (**C**) Example microhabitats: a city park (*pipiens)* and the flooded basement of an apartment complex (*molestus*). (**D**) Two hypotheses describing *molestus*’ origin. Hypothesis 1 (left) posits that belowground *molestus* evolved from local aboveground *pipiens* in situ within the past 100–200 years. Hypothesis 2 (right) posits that *molestus* first evolved in an aboveground context thousands of years ago, possibly in association with early agricultural societies of the Mediterranean basin, with colonization of belowground habitats (dotted arrow) occurring much later (*22*, *23*). [Image credits: Lawrence Reeves (mosquito); Yuki Haba (city park); Colin Malcolm (flooded basement)]

The sophisticated adaptations of *molestus* to urban belowground environments have led to much speculation over when and where it originated. A widely cited hypothesis suggests that *molestus* evolved in the London Underground subway system, where it first became famous in the 1940s during World War II (*1*, *6*–*11*). At that time, many Londoners took nightly refuge in the city’s subway system to escape intense Nazi bombing. Sleeping on subway platforms protected people from bombs but made them easy targets for *molestus*, which became known as the ‘London Underground Mosquito’ and was hypothesized to have evolved there during the ∼100 year period between subway tunnel construction and mosquito discovery (Fig. 1D, left) (*6*, *12*). Form *molestus* was reported in cellars and cesspits in France, Denmark, Germany, and the former USSR 10–25 years before its discovery in London (*3*,*13*–*15*), but an urban, belowground origin in northern Europe within the past few hundred years remains possible. Recent reviews have pointed to *molestus* as one of the best candidates for rapid urban adaptation (*1*, *7*–*11*), and major science news outlets treat this hypothesis as fact (*16*–*21*). The idea that such an array of traits could emerge *de novo* in just a few hundred years is striking and sets a new bar for the number and complexity of changes we might expect to occur in modern cities over short timescales.

An alternative hypothesis, which is mentioned but less prominent in the literature, posits that *molestus* first adapted to humans in an aboveground context, long before the rise of modern cities (Fig. 1D, right) (*22*, *23*). While *molestus* is confined to belowground habitats in cold regions, it thrives aboveground in warmer climates, particularly in the Mediterranean basin (*23*). Moreover, early records document *molestus*-like mosquitoes breeding and biting humans aboveground in Egypt, Croatia, and Italy 50–100 years before they were discovered in basements and subways (*24*–*26*). According to this alternative scenario, many of the traits that allow *molestus* to thrive in urban belowground environments would represent exaptations, or traits that first arose in a different time and context (*27*). An aboveground Mediterranean origin could also push the timing of *molestus*’ origin back thousands of years, to an era when humans first started forming dense agricultural communities. Early allozyme and microsatellite studies indicate that contemporary *molestus* populations from aboveground and belowground habitats are genetically related (*22*, *28*), but the validity and timing of a putative aboveground origin remain to be tested.

Here we leverage the first large population genomic dataset for *Cx. pipiens* to infer when, where, and in what ecological context *molestus* first evolved. Beyond its enigmatic origins, *molestus* is a competent disease vector, implicated in the transmission of West Nile virus and other arboviruses across Eurasia and North America over the past several decades (*29*, *30*). Solving the mystery of *molestus*’ origins thus has important implications for understanding both rapid urban adaptation and emerging threats to human health.

### Form *molestus* is genetically isolated from *pipiens* across the Western Palearctic

Multiple lines of evidence indicate that *molestus* first split from *pipiens* somewhere in the Western Palearctic (a region that includes Europe, North Africa, and western Asia) (*31*), before spreading to other parts of the world. However, the structure of populations across this region has been difficult to decipher due to the absence of morphological differences. Analysis of one or a small number of genetic loci shows that the two forms are isolated in northern Europe, where harsh winters confine *molestus* to belowground environments (*22*, *23*). However, *molestus* and *pipiens* appear to be more genetically similar in southern Europe, where both breed aboveground, and may even collapse into a single panmictic population in North Africa (*22*, *23*). To better resolve the situation with high-resolution genomic data, we sequenced the whole genomes of 357 *Cx. pipiens* individuals collected in 77 locations scattered across the Western Palearctic (Fig. 2A; *n* = ∼5 individuals per population at 12.9X median coverage). These data are part of a larger collection of 840 genomes to be presented in a companion study of the deeper evolutionary history of *Cx. pipiens* across its entire global range (*32*).

**Fig. 2.**
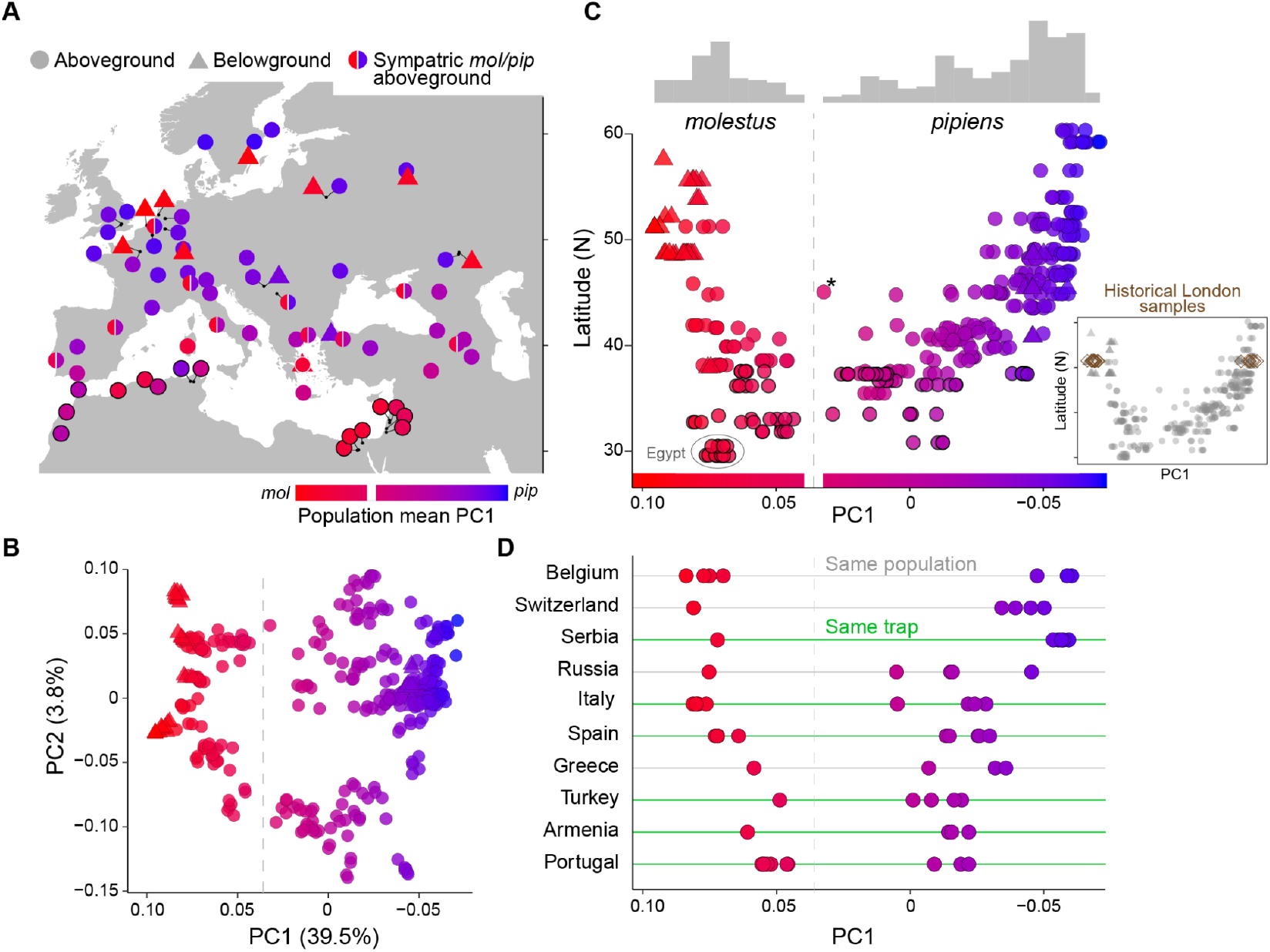
Form *molestus* is genetically isolated from *pipiens* across the Western Palearctic. (**A**) Sampled populations, colored by average PC1 value. Circles and triangles represent aboveground and belowground locations, respectively. Half-circles indicate that both *pipiens* and *molestus* were collected in the same or nearby aboveground sites. (**B**) PCA of genetic variation across all samples in (A) (*n* = 357). (**C**) PC1 values plotted against latitude, with marginal frequency histogram at top. The gray dashed line indicates a natural break in the histogram, inferred to separate *pipiens* and *molestus* (PC1 = 0.04). Thick outlines mark individuals from North Africa and the Middle East. Asterisk marks a putative F1 hybrid from southwest Russia (Stavropol). Inset shows position of historical London samples in a combined PCA with contemporary mosquitoes (*n*=22, collected 1940-1985; see also fig. S1). (**D**) PC1 values for *pipiens* and *molestus* individuals collected in the exact same day and trap (green lines) or in the same general location (within 5–45 km; grey lines).

We used a Principal Component Analysis (PCA) to assess variation across the Western Palearctic using 504,000 high quality SNPs (*32*) (Fig. 2B). The first major axis (PC1) accounted for by far the most variation (39.5%, fig. S2A) and was thus likely to represent divergence between *pipiens* and *molestus*. Consistent with this prediction, belowground and aboveground samples from northern latitudes were clustered at opposite ends of the PC1 axis (Fig. 2C). Sequenced mosquitoes with known biting or egg-laying behavior (*n* = 13), including those from lower latitudes, were also arrayed across PC1 according to expected form (fig. S1A). PC2 explained ∼4% of genetic variation across the sample (Fig. 2B) and was strongly correlated with longitude (fig. S2B).

Our sample included aboveground mosquitoes from London, which clustered with other northern European *pipiens*, but we were not given permission to collect mosquitoes in the London Underground. To confirm that the genetic picture today reflects the one present when iconic WWII populations were first discovered, we used a minimally destructive approach (*33*) to extract and sequence DNA from 22 museum specimens collected at 15 sites in London between 1940 and 1985 (table S2, mean genome-wide coverage = 5.8X). Metadata for most samples did not specify microhabitat, but the sampling locations included the sites of major underground stations, including Paddington, Monument, and Barking. A joint PCA with contemporary samples placed the historical London specimens in the same two genetic clusters that characterize mosquitoes at that latitude today (Fig. 2C, inset; fig. S1B). We conclude that the genetic character of *pipiens* and *molestus* populations in northern Europe has been stable for the past 75 years.

Form *pipiens* and *molestus* are genetically well separated in the north, but our data confirm that they are less distinct at southern latitudes. Mosquitoes on both the *molestus* and *pipiens* ends of the PC1 axis have increasingly intermediate values as one moves from northern Europe towards Africa, creating a striking U-shaped pattern when PC1 is plotted against latitude (Fig. 2C). Importantly, however, they never completely merge; even southern populations fall into two discrete genetic clusters with a break at PC1 ∼0.04 (Fig. 2C, dashed line). Moreover, individuals from these two clusters were frequently collected in the same traps, highlighting the absence of microgeographic barriers (Fig. 2D). Our whole genome data thus show unequivocally that *pipiens* and *molestus* are able to coexist in sympatry across the region (*34*, *35*). They are genetically closer in the south, and several individuals in our sample may represent early generation hybrids (e.g. see asterisk in Fig. 2C). Despite this, we see no evidence of collapse into a panmictic population.

### Ancestral latitudinal gradient within *pipiens* suggests *molestus* arose at the southern edge of the Western Palearctic

The genetic similarity of *molestus* and *pipiens* at southern latitudes is believed to result from increased gene flow (*22*, *23*); hybridization should be rare in the north where the two forms occupy different microhabitats, but increasingly common in the south where both breed aboveground (Fig. 3A). To test this hypothesis we examined the latitudinal cline within *pipiens*, which is much stronger than that within *molestus* (Fig. 2C). More specifically, we used genome-wide *f3* statistics (*36*) to model each *pipiens* population as a mix of ‘pure’ *pipiens* and *molestus* reference populations taken from their northern extremes. Many European and west Asian populations showed evidence of mixing (Fig. 3B), but the signal was not latitudinal (Fig. 3C; Pearson’s *r* = −0.022, *P* = 0.90). Moreover, North African *pipiens* populations, which are genetically closest to *molestus*, showed no signs of admixture (Fig. 3B). These results cast doubt on the longstanding hypothesis that latitudinal variation within *pipiens* is driven by hybridization with *molestus*.

**Fig. 3.**
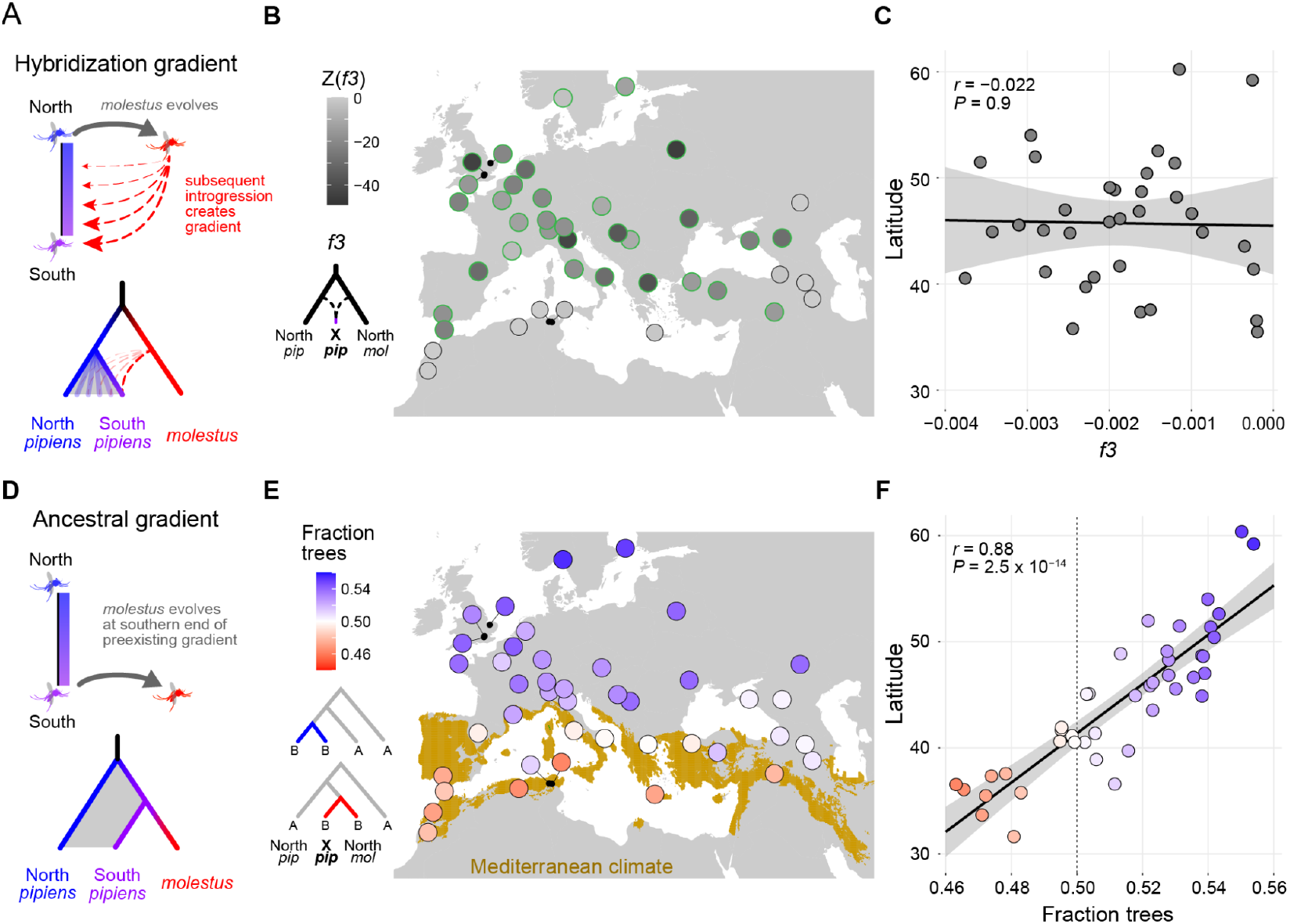
Ancestral latitudinal gradient within *pipiens* suggests *molestus* arose at the southern edge of the Western Palearctic. (**A**) Hybridization gradient hypothesis: the genetic gradient within *pipiens* may result from increasing levels of gene flow with *molestus* as one moves from north to south. (**B**) *Z* scores of genome-wide *f3* values for each *pipiens* population when modeled as a mixture of northern *pipiens* (Sweden) and northern *molestus* (Belgium). Significantly negative *f3* values (*Z* < −3, green outlines) are consistent with the presence of admixture. (**C**) *f3* statistics of *pipiens* populations with significant signs of admixture, plotted against latitude. (**D**) Ancestral gradient hypothesis: the genetic gradient within *pipiens* may be ancestral, with *molestus* evolving from southern *pipiens* populations. (**E**) Fraction of derived alleles shared by each *pipiens* population with northern *pipiens* versus northern *molestus. Culex torrentium* was used as the outgroup. Light brown in the map shows the Mediterranean climate zone. (**F**) Fraction of derived alleles shared with northern *pipiens* vs. *molestus*, plotted against latitude. Both (C) and (F) include linear regression line with 95% confidence interval and Pearson’s correlation test statistics. Across all analyses, only populations with four or more individuals were included.

An alternative hypothesis, which has not been explored in the literature, is that the latitudinal gradient within *pipiens* predates the evolution of *molestus*. In this case, southern *pipiens* could be genetically closer to *molestus* not because they mix with *molestus*, but because they gave rise to *molestus* (Fig. 3D). Consistent with this idea, we found that southern *pipiens*—and especially *pipiens* populations in the Mediterranean basin—share as many, or more, derived alleles with a reference *molestus* population from the north than they do with a reference *pipiens* population from the north (Fig. 3E). Moreover, the overall signal of relative allele sharing was strongly latitudinal (Fig. 3F, Pearson’s *r* = 0.88, *P* = 2.5 x 10^-14^). Taken together, we conclude that the latitudinal gradient within *pipiens* is ancestral—perhaps reflecting adaptation to variation in temperature and/or precipitation—and that *molestus* is most likely derived from populations in the south.

### Form *molestus* evolved thousands of years ago in the Middle East

We further explored the geography of *molestus’* origin by constructing a distance-based (*Dxy*) tree for *Cx. pipiens* individuals from the full global sample (*32*). Contemporary gene flow can obscure ancestral relationships in phylogenetic trees. We were therefore careful to exclude any individual or population that showed signs of recent introgression from the other form (fig. S3–S4, S8) or from *Culex quinquefasciatus*, a tropical sibling species that hybridizes with *Cx. pipiens* in the Americas and Asia (*32*). We also excluded low coverage samples (<10X), leaving a total of 205 individuals.

The resulting tree provided strong support for a southern origin of *molestus*. First, all *molestus* samples formed a monophyletic clade that was nested within Mediterranean *pipiens* (Fig. 4A, fig. S5). Second, the earliest branching lineages within *molestus* corresponded to aboveground mosquitoes from the eastern Mediterranean—specifically Egypt, Israel, and Greece (Fig. 4A). Egyptian and Israeli samples were also among the most genetically diverse, together with two populations from the Caucasus region (Armenia and southern Russia) (Fig. 4B). Finally, while belowground *molestus* from northern latitudes formed tight, derived clades, aboveground *molestus* populations from North Africa, the Middle East, and southern Europe were scattered across the base of the tree (Fig. 4A). These results support the hypothesis that *molestus* first evolved in an aboveground context in the greater Mediterranean basin, and more specifically in the Middle East.

**Fig. 4.**
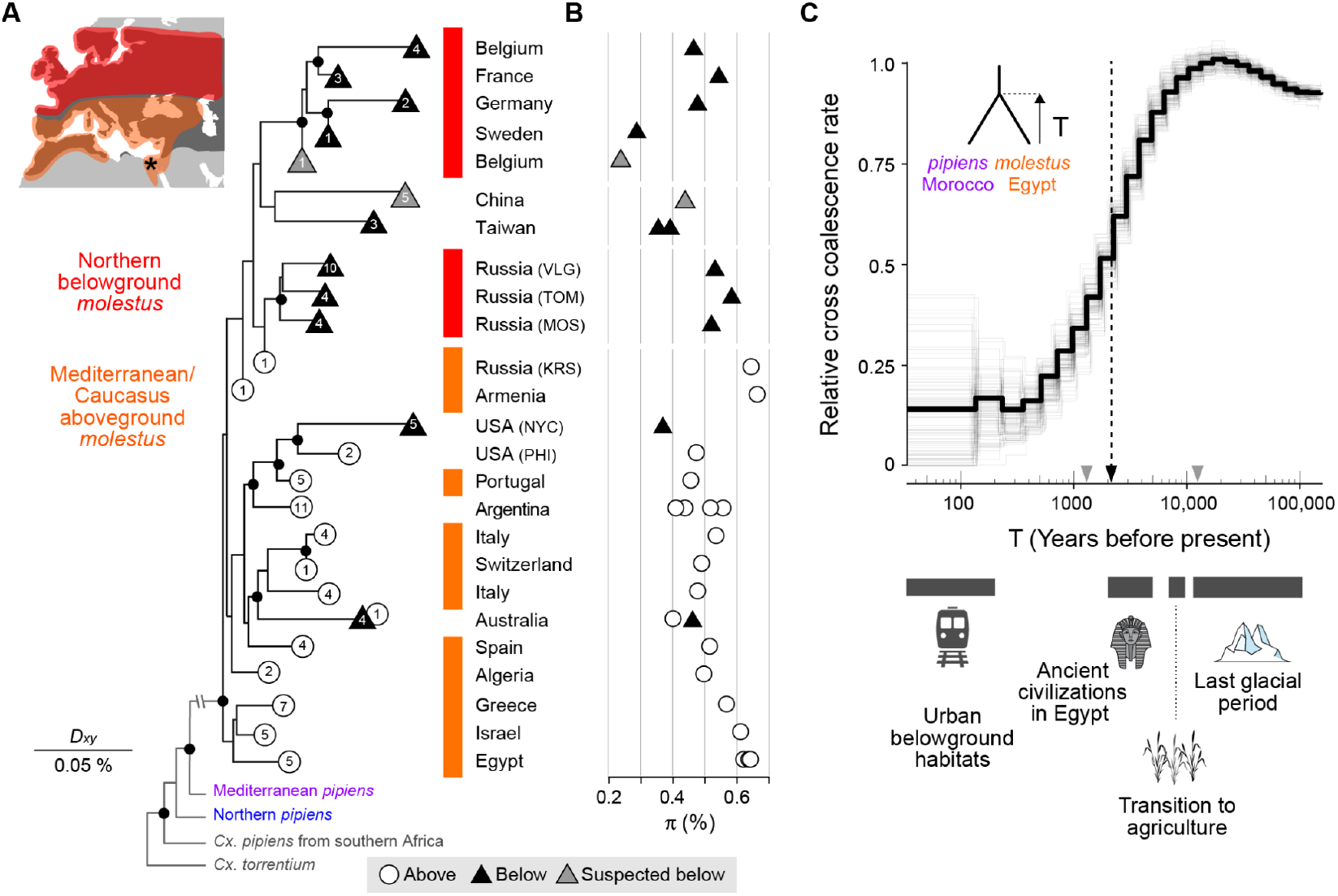
Form *molestus* evolved thousands of years ago in the Middle East. (**A**) *molestus* clade excerpted from neighbor-joining tree based on pairwise genetic distance (*Dxy*) among putatively unadmixed *pipiens* and *molestus* individuals from the global sample (*32*). Terminal branches are collapsed at the root of each population, with a symbol and number indicating microhabitat and sample size, respectively (*32*). Map inset shows distribution of two subgroups of *molestus* from the tree (orange/red) and *pipiens* (dark grey). Only *molestus* is present in Egypt, marked by an asterisk. Black circles mark nodes with >95% bootstrap support. See fig. S5 for the full tree. (**B**) Genome-wide nucleotide diversity (π) of populations shown in (A). (**C**) Relative cross-coalescence (rCC) rate between Moroccan *pipiens* (MAK) and Egyptian *molestus* (ADR) inferred from phased, whole genome sequences (*32*). rCC rate is expected to plateau at 1, going backwards in time, when populations have merged into a single ancestral population. Rapid divergence is observed between ∼10K and 1K years ago, with a split time (T; rCC rate=50%) of 2,141 years (black dashed arrow) based on estimates of generation time and mutation rate (*32*). Biologically reasonable upper and lower bounds for these parameters give minimum and maximum split times of 1,298 and 12,468 years (grey arrowheads). Thick black line shows genome-wide result, and light grey lines show 100 bootstrap replicates. Illustration credits: Wheat, Freepik.com. Pharaoh and Glacier, Vecteezy.com.

The Middle East is a compelling location for the emergence of *molestus* not only because it harbors diverse, early branching populations (Fig. 4A–B), but also because it is the only place within the Western Palearctic where *molestus* is known to occur on its own, in the absence of *pipiens* (Fig. 2A) (*28*, *37*). The Middle East was also home to some of the earliest agricultural societies, which were thriving in Mesopotamia and Egypt by 3000 BCE (*38*). A Middle Eastern origin thus raises the possibility that *molestus* first adapted to human hosts and habitats in isolation from *pipiens* and on a timescale of thousands, rather than hundreds, of years (Fig. 1D, right) (*22*, *23*).

We explored the timing of *molestus*’ origin using a cross-coalescent analysis of DNA haplotypes from Middle Eastern *molestus* (Egypt) and Mediterranean *pipiens* (Morocco) (*n* = 2 individuals with ∼50X coverage from each population) (*39*). As expected, the relative cross coalescence (rCC) rate starts near zero in the recent past, when the two populations are isolated, but rises monotonically and eventually plateaus near one, going backward in time, when they merge into a single ancestral population (Fig. 4C). Accurate assignment of dates to this rCC curve requires knowledge of the *de novo* mutation rate (*μ*) and generation time (*g*), neither of which has been directly measured for *Cx. pipiens* in nature. However, plausible literature estimates for *μ* (4.85×10^−9^) and *g* (20 days) (*32*) suggest that peak rates of divergence occurred ∼2,000 years ago (Fig. 4C, dashed arrow), while minimum and maximum reasonable values (*32*) lead to split times anywhere between 1,300 and 12,500 years ago (Fig. 4C, grey arrowheads). We obtained a similar range of split times when using an alternative *pipiens* population from the southern Caucasus region (Armenia; fig. S7). Taken together, these results are inconsistent with a post-industrial origin for *molestus* in northern Europe (Fig. 1D, left) and instead support an ancient origin associated with early agricultural civilizations of the Middle East (Fig. 1D, right).

### Introgression from *molestus* into aboveground *pipiens* is associated with human density

Recent urbanization did not drive initial evolution of *molestus*, but it may have driven its expansion and increased contact with *pipiens* across the northern hemisphere—contact that is thought to have contributed to the emergence of West Nile virus (WNV) in human populations over the last several decades (*29*, *30*). WNV is a mosquito-borne virus that primarily infects birds and is effectively amplified within avian populations by bird-biting *pipiens* (Fig. 5A). Spillover to dead-end human hosts can only occur if local *pipiens* mosquitoes are also willing to bite humans, a broadening of biting behavior that may be driven by gene flow from *molestus* in urban areas (*40*–*42*). This idea has spurred efforts to detect and quantify admixture between *pipiens* and *molestus* in natural populations (*23*), yet we show above that much of the genetic signal previously attributed to mixing between forms instead represents ancestral variation (Fig. 3). Form *pipiens* likely receives genetic input from *molestus* in some places, but exactly where and to what extent are not known.

**Fig. 5.**
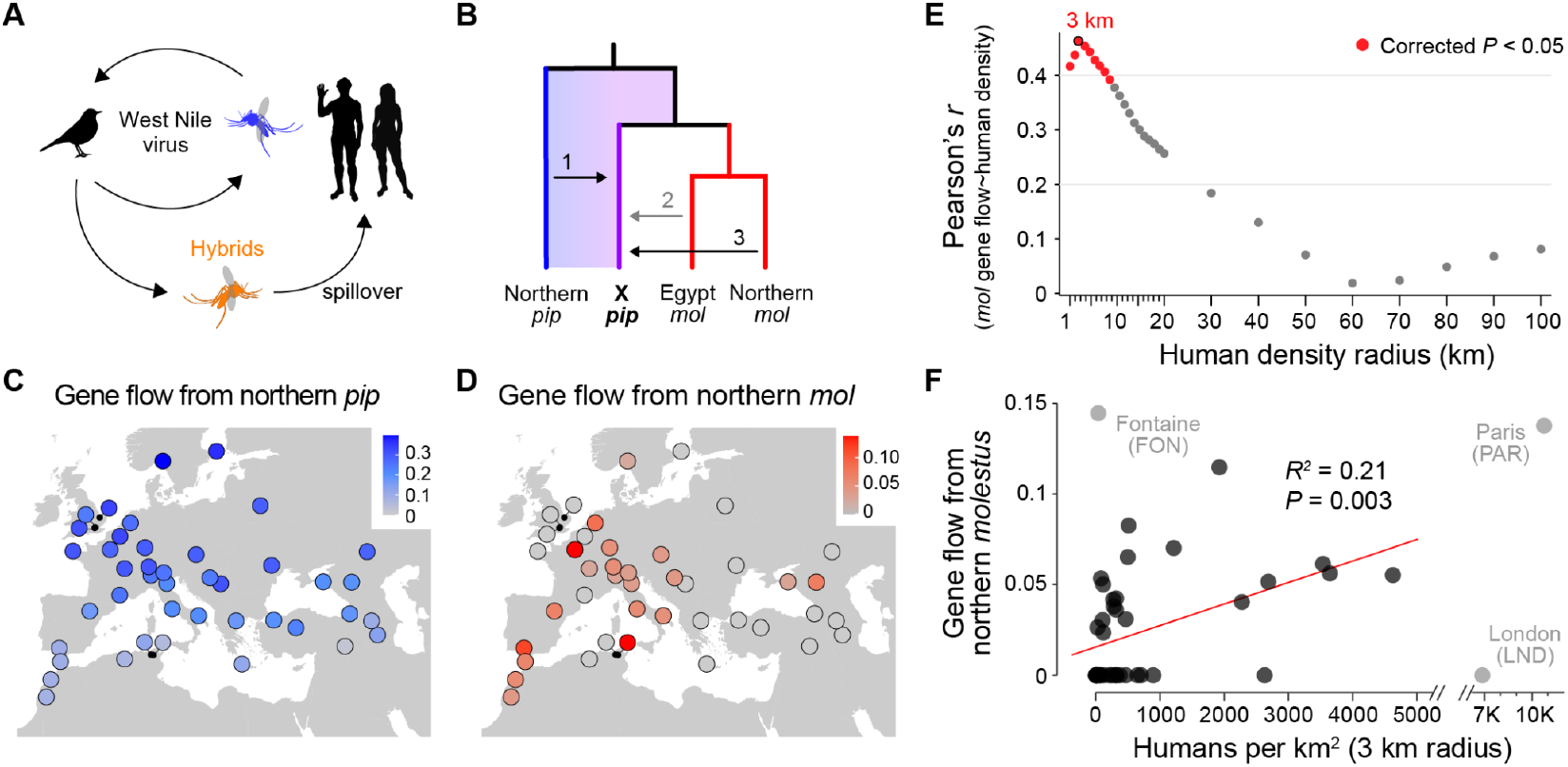
Introgression from *molestus* into *pipiens* is associated with human density. (**A**) Schematic of West Nile virus transmission dynamics. Form *pipiens*-*molestus* hybrids, which have intermediate biting preference (*42*), are implicated in the spillover of WNV from birds to humans (*40*, *41*). (**B**) Base tree used to simultaneously estimate three potential sources of introgression into focal *pipiens* populations (X *pip*): northern *pipiens* (Sweden, SWE), Middle Eastern *molestus* (Egypt, ADR), and northern *molestus* (Belgium, BVR). (**C**–**D**) Population-specific estimates of ‘gene flow’ from northern *pipiens* (C), which accounts for the ancestral latitudinal gradient, and northern *molestus* (D). We did not detect gene flow into any population from Middle Eastern *molestus* (fig. S8). (**E**) Correlation between gene flow from northern *molestus* (D) and human population density within circles of varying radius around each collection site. (**F**) Gene flow from northern *molestus* as a function of human density within a 3-km radius of each collection site. Three outliers in grey (Cook’s distance > 4) were excluded, but regression remains significant if included (*P* = 0.005, *R*^*2*^ = 0.18). [Image credit: Phylopic.com (bird and human)]

We used *f*-branch statistics to reassess levels of gene flow from *molestus* into *pipiens* while controlling for ancestral variation. *f*-branch statistics allow simultaneous quantification of gene flow among multiple branches in a tree (*43*). To account for the sister relationship between forms in the south (Fig. 3D), we specified a fixed tree in which focal *pipiens* populations were genetically closer to *molestus* than to a reference *pipiens* population from the far north (Fig 5B). Deviations from this topology (i.e. for focal populations from the north) can then be modeled as significant ‘gene flow’ into the focal *pipiens* population from the northern reference (Fig. 5B, arrow 1). As expected, the resulting signal was strongly latitudinal (Fig. 5C, fig. S8A). The tree also included two potential sources of *molestus* introgression, allowing us to distinguish them. We could not detect gene flow into any *pipiens* population from the early branching *molestus* lineage in Egypt (Fig. 5B arrow 2; fig. S8C), consistent with its isolated location at the southern edge of the contemporary range. However, we detected substantial gene flow into some *pipiens* populations from a derived *molestus* lineage present in the north (Fig. 5B arrow 3, Fig. 5D, fig. S8B).

Levels of gene flow from northern *molestus* into *pipiens* did not covary with latitude (Fig. 5D, fig. S8B), but showed a positive association with human population density (*44*–*46*). The more humans living within 1–10 km of each sampling location, the more likely we were to observe introgression (Fig. 5E). This relationship was most significant when averaging human density across an area with 3 km radius (linear regression *P* = 0.003, *R*^*2*^ = 0.21, Fig. 5E–F), suggesting that levels of urbanization immediately around collection sites are most predictive of hybridization. Moreover, this signal was driven primarily by the consistent presence of ∼5% introgression in truly ‘urban’ areas, defined by the European Commission as having >1500 people per km^2^ (Fig. 5F) (*47*). Introgression was less predictable at rural sites (*P* = 0.07, *R*^*2*^ = 0.11 excluding urban centers). Inclusion of three statistical outliers in this analysis (Fig. 5F, grey dots) slightly weakened the trend, but still indicated a strong association (regression *P* = 0.005, *R*^*2*^ = 0.18). Interestingly, a site in Paris showed ∼15% introgression, as one might expect based on its density, but we could not detect any genetic input from *molestus* in London. Such geographic variability in levels of gene flow may in part reflect whether local *pipiens* and *molestus* populations are infected by compatible or incompatible strains of *Wolbachia pipientis* bacteria (*48*, *49*). Taken together, our results counter the longstanding idea that gene flow between *pipiens* and *molestus* is greatest at southern latitudes (where both forms breed aboveground) and instead reveal a complex landscape of introgression that is modestly associated with levels of human activity.

## Discussion

Understanding how life can adapt to rapid urbanization is an important challenge in evolutionary biology. As examples accumulate in the literature, each new case provides a reference for the potential speed and character of adaptation. Here, we revisit one of the most iconic such examples using high resolution population genomic data. Instead of evolving in the subway system of a northern European city over the course of 100–200 years, our results suggest that *Cx. pipiens* f. *molestus* first adapted to human hosts and habitats in the Middle East over the course of 1000 or more years, possibly in association with early agricultural societies (Fig. 6). Early agricultural settlements would have provided a novel ‘human’ niche in a region that might otherwise have been too arid to support robust *Cx. pipiens* populations. Irrigation systems offer rich breeding sites for larval stages, while abundant humans and domestic animals offer a reliable source of blood for adult females. We cannot say where exactly within the Middle East adaptation first occurred, as our sampling in the region is limited and ranges shift over time. However, *molestus* is particularly abundant in Egypt’s Nile basin, and ancient pharaonic artifacts and papyrus are consistent with the idea that *molestus* was spreading filarial worms among humans there as many as 2000 years ago (*23*).

**Fig. 6.**
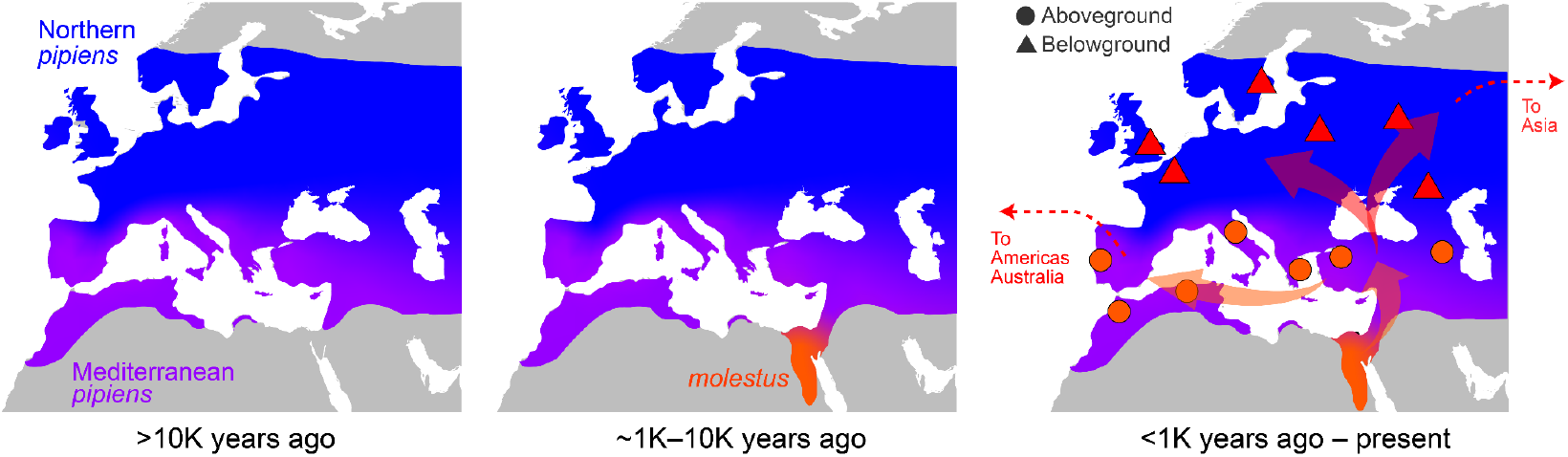
Inferred evolutionary history of *molestus*. Three sequential panels show the inferred history of *molestus*. **Left**: The latitudinal genetic gradient characterizing extant *pipiens* populations in the Western Palearctic predates *molestus*. **Middle**: The rise of dense, settled, agricultural communities in the Middle East 2000–10,000 years ago, including along the Nile River in Egypt, would have provided new ecological opportunities for mosquitoes that could adapt to human hosts and habitats—driving the evolution of *molestus*. **Right**: Eventually *molestus* must have spread north, where it became established alongside *pipiens* in the warm Mediterranean region by the 1800s (but possibly earlier) and in belowground microhabitats of cold, northern cities by the early 1900s. Our distance tree (Fig. 4A) suggests that *molestus* was further spread (most likely by humans) overland all the way to East Asia and from the Mediterranean region overseas to America and Australia.

Rather than benchmarking the speed and complexity of urban evolution, this updated history highlights the role of preexisting traits, or exaptations, in adaptation to urban environments (*27*). Three of the key behaviors that allow *molestus* to thrive belowground are present in contemporary Egyptian populations and almost certainly arose in ancient times: mammal-biting, the ability to mate in confined spaces, and the ability to lay a first clutch of eggs without a blood meal (*37*, *50*). A fourth trait, lack of diapause, which limits *molestus* to belowground environments at northern latitudes, is also present in the Middle East today (*37*). Form *molestus* was thus primed to take advantage of northern, belowground environments before they arose. Indeed, it joins a host of other ‘urban’ taxa that first became dependent on humans thousands of years ago, including brown rats (*51*), house mice (*52*), cockroaches (*53*), house sparrows (*54*), and the dengue mosquito *Aedes aegypti* (*55*).

Ancient origins do not preclude additional, contemporary evolution (*51*, *53*). Once established belowground, *molestus* was likely exposed to a new suite of challenges. For example, many belowground habitats lack vertebrate hosts altogether, providing a competitive edge to females that can develop eggs without a bloodmeal. This trait, called autogeny, is present in contemporary Middle Eastern *molestus*, but only at low frequency (*37*, *50*). In contrast, it occurs at high frequency in some aboveground *molestus* from southern Europe (*34*) and is nearly fixed in northern belowground populations (Fig. 1B). Future work should explore whether increased autogeny may provide a bona fide example of rapid, urban evolution in belowground environments and whether this change arose just once, or many times in parallel (*56*). Our distance tree (Fig. 4A) suggests that belowground populations from northern Eurasia are all closely related, but that those from the east coast of North America (and possibly Australia) represent independent colonization events.

Our findings also carry public health implications. The emergence and spread of WNV over the last two decades has triggered an intense interest in quantifying admixture between *molestus* and *pipiens*, as hybridization is thought to drive spillover from birds to humans (*40*, *41*). Yet we show that true patterns of admixture are obscured by ancestral relationships at southern latitudes. In particular, it is currently standard practice to identify ‘hybrids’ using a single locus marker called CQ11 (*57*). Pure *pipiens* and *molestus* were thought to be fixed for different alleles at this locus such that heterozygotes must be ‘hybrids’. Our data instead suggest that *pipiens* harbors ancestral variation at this locus. The ‘*molestus*’ allele was likely present at moderate frequency in the Mediterranean basin before *molestus* arose, and remains present in ‘pure’ Mediterranean *pipiens* today. Accurate inferences of gene flow will require more substantial genomic data and relatively complex analytical methods (e.g. Fig. 5). Future work should also consider the possibility that Mediterranean *pipiens* populations are somewhat intermediate between canonical northern forms at the phenotypic, as well as genetic, level (*58*). They may be effective bridge vectors even in the absence of genetic input from *molestus*. WNV represents an increasing threat to public health across the northern hemisphere, with many of the most severe outbreaks occurring within the last 5 years (*59*, *60*). Taken together, we hope that our work opens the door to more incisive investigation of the potential links between urbanization, gene flow, ancestral variation, and viral spillover.

## Acknowledgements

We thank members of the McBride Lab for discussion.

## Author Contribution

YH conceived the study, performed and analyzed experiments, and wrote the manuscript. PK, EM, and MKNL helped conceive, collect, and analyze data associated with archival samples from London. MS helped advise YH and analyze experiments. NHR helped conceive the study and analyze experiments. CSM conceived and supervised the study, helped analyze experiments, and wrote the manuscript. All other authors collected and identified mosquitoes. All authors provided feedback on the manuscript.

## Data and materials availability

All genome resequencing data associated with this study is available in the NCBI SRA (790 contemporary genomes PRJNA1209100, 22 genomes from archival museum specimens ERS10924505–ERS10924526). Associated scripts will be available on GitHub (github.com/YukiHaba).

## Supplementary Materials

Materials and Methods Figs. S1 to S8

Tables S1 and S2

References (*61*)*-*(*93*)

## Materials and Methods

### *Culex pipiens* Population Genomics Project

This study is one of two flagship studies associated with the *Culex* pip*iens* Population Genomic Project, also known as PipPop. Both studies make use of 840 individual whole-genome sequences of *Culex pipiens* complex mosquitoes (*Cx. pipiens sensu lato*) and outgroups (Table S1). Within the complex, we specifically targeted *Cx. pipiens s. s*. Linnaeus, 1758 and hybrids (*n*=688), but also sequenced smaller numbers of *Cx. quinquefasciatus* Say, 1823 (*n*=101), *Cx. pallens* Coquillett, 1898 (*n*=33), and *Cx. australicus* Dobrotworsky & Drummond, 1953 (*n*=5). *Cx. torrentium* Martini, 1925 (*n*=9) was included as an outgroup, and a handful of sequenced mosquitoes were inferred to belong to more distant, unknown taxa (*n*=4; Table S1). A total of 790 genomes were sequenced for PipPop, while 50 were previously published (40 from (*61*) and 10 from (*62*)). Full details on sampling of the 790 PipPop genomes, as well as variant calling and sample filtering for the full dataset, are provided in the companion study and summarized briefly here.

#### Mosquito collection and sequencing

We collected and sequenced 790 mosquitoes from 163 populations spread across 44 countries in the Americas, Europe, Africa, Asia, and Australia, targeting *n* ∼ 5 individuals per population. 752 mosquitoes (95%) were collected from 2014–2021, and the remaining 38 (5%) were collected from 2003–2012. Mosquitoes were mostly sampled as adults and identified as *Cx. pipiens s. l*. or *Cx. torrentium* using standard metrics, but a subset came from dense larval pools. They were collected from both aboveground (87%) and belowground (13%) sites. Belowground sites included basements of residential buildings, manholes, stormwater drains, cesspits, subway systems, and underground floors of a parking garage. Aboveground sites spanned a variety of habitats, from dense urban environments to residential areas to natural parks. Detailed sample metadata, including individual and population IDs, GPS coordinates, collection date, life stage, sex, and trapping method can be found in Table S1.

Genomic DNA was extracted using the NucleoSpin 96 DNA RapidLyse kit (Macherey-Nagel, Germany). We confirmed that samples belonged to the *Cx. pipiens* complex or *Cx. torrentium* (outgroup) via a multiplex PCR targeting the *ace-2* locus (*63*) and visual inspection of amplicon sizes on a gel. Samples with no bands or unexpected band sizes were excluded. DNA sequencing libraries were prepared using Illumina DNA Prep Kits (Illumina, USA) with custom dual-unique barcodes. Approximately 80 barcoded libraries were pooled and sequenced on individual S4 lanes of a Novaseq 6000 PE150 sequencer (Illumina, USA), with a target genome-wide coverage of 10-15X. One pool including Mediterranean and Middle Eastern mosquitoes was sequenced across four lanes (a full S4 flow cell) to achieve higher coverage (∼60X) for use in cross-coalescence analyses.

#### Read processing and mapping

Raw reads were assessed for quality using FastQC v.0.11.8 (*64*), and low-quality bases and adapters were trimmed using Trimmomatic (*65*). Trimmed reads were mapped onto the recently updated, chromosome-scale CpipJ5 assembly (*62*). We used BWA-MEM v.0.7.17 (*66*) to map the reads with default settings and identified and removed optical and PCR duplicates with Picard MarkDuplicates v.2.20.2 (*67*). We then used GATK v.3.8 (*68*) to perform local realignment around small insertions and deletions. We calculated genome-wide coverage after deduplication using Mosdepth v.0.3.3 (*69*). We used the deduplicated, realigned reads for all the analyses below.

#### Accessible regions and variant calling

We used 100 high-coverage individuals (50 *Cx. pipiens s. s*. with >20X coverage and 50 *Cx. quinquefasciatus* with >10X coverage) to characterize variation in coverage across the genome and mask ‘inaccessible’ regions for variant calling. More specifically, we masked sites with <0.5X or >1.5X normalized coverage in either species. We also masked putative repeat elements regardless of coverage (*62*). This left us with ∼131 million ‘accessible’ sites or approximately 23% of the 559 Mb genome. We then called single nucleotide variants in all 840 individuals using BCFtools v1.13 (*70*). Variant calling was parallelized across multiple 20 Mb chunks of the genome. In addition to masking the inaccessible sites and repeat elements described above, we also masked multiallelic SNPs, indels, and SNPs falling within 5 bp of indels. We calculated key statistics for each SNP and further removed those with QUAL<50, MQ<50, >10% individuals with missing genotypes, average mean depth across all samples of <10X or >30X, and alleles of GQ<20. These cutoffs were chosen after visual inspection of the distribution of each statistic, following GATK hard-filtering best practices for non-model species (*71*). After filtering, we were left with 30.6M high-quality, accessible, biallelic SNPs (of ∼131M total accessible sites). This full SNP set was used for all analyses except where otherwise specified.

#### Individual filtering

We removed two samples from Raleigh, USA with < 2X coverage and >50% genotype missingness (RAL5, RAL6). We also filtered the full sample set for kin based on pairwise KING kinship coefficients computed in NgsRelate v.2.0 (*72*). The vast majority of pairs showed low relatedness as expected (mean kinship coefficient 0.00026). However, a subset of pairs showed higher values, including a subset of mosquitoes collected as larvae in the same pools. We identified all pairs with kinship > 0.09 and excluded the individual with lower coverage. We additionally excluded 3 individuals (PAR4, OSJi4, KAV5) that showed unexpectedly high relatedness to many individuals from other populations and one individual from Malaysia (MEL5) that clustered with North American samples. The unexpected relatedness of PAR4, OSJi4, and KAV5 to many other individuals could not be explained by their position in the 96-well plates used to process samples nor by low sequence coverage. While the Malaysian sample could conceivably be a migrant, we chose to remove it out of an abundance of caution.

#### Final sample set

After filtering low quality individuals and kin, we were left with data for 743 unrelated mosquitoes. A few analyses presented here address the full global sample. However, unless otherwise specified, this study focuses on the subset of 357 individuals collected in the Western Palearctic (Europe, North Africa, and western Asia).

### Analysis of population structure

We conducted a principal component analysis (PCA) of variation among Western Palearctic individuals (*n* = 357; Fig. 2 and fig. S1). As excessive linkage disequilibrium (LD) among genetic markers can lead to PC(s) of LD structure rather than population structure (*73*), we used Plink v.1.90 (*74*) to select a subset of 503,921 unlinked SNPs (--indep-pairwise 200 20 0.2). We then used PCAngsd v.1.10 (*75*) to estimate a covariance matrix and the `princomp` function in the R package `stats` v.3.6.2 to conduct the PCA (*76*).

### Sequencing and analysis of historical specimens from London

To understand the relationship between historical and contemporary *molestus* populations, we extracted genomic DNA from 22 pinned *Culex* specimens in the National History Museum, London (Table S2) using a recently published, minimally destructive protocol (*33*). Briefly, pinned specimens were removed from the main label pins and put in a styrofoam box filled with wet paper towels for rehydration at 37 °C for 3 hours. Each rehydrated sample was then dipped in 200 µl of Lysis Buffer C (200 mM Tris, 25 mM EDTA, 0.05% Tween-20, and 0.4 mg/ml Proteinase K) and incubated at 37 °C for 2 hours. Genomic DNA in the lysis buffer was then purified using a modified MinElute (Qiagen) silica column approach. After extraction, intact mosquito specimens were rinsed in increasing percentages of ethanol (30% and 50%) and sent back to the museum for critical point drying. Libraries of the purified genomic DNA were created using NEB Next Ultra II DNA Library Prep Kit (New England Biolabs) with no shearing and then purified using 2.2x SPRI (Beckman Coulter Agencourt AMPure XP) beads post library ligation and two times 1x SPRI post PCR amplification using a KAPA HiFi HotStart Uracil+ ReadyMix PCR Kit. The final libraries were sequenced on one lane of NovaSeq PE75 (Illumina).

Raw reads were run though the ancient DNA pipeline EAGER (*77*), with the following processing parameters: trimming adapter sequence, trimming bases of quality score < 20, removing sequences shorter than 30 bp, merging overlapping paired reads (with default minimum 11 bp overlap), aligning to the CpipJ5 assembly (*62*) using BWA-MEM, removing PCR duplicates and unaligned reads for final BAM files, and performing DamageProfiler to summarize ancient DNA characteristics (50 C > T and 30 G > A substitutions, read length in base pairs). We calculated genome-wide coverage after deduplication using Mosdepth (*69*) (mean = 5.77X, range = 1.05–9.22X). We used ANGSD v.0.936 (*78*) to call genotype likelihoods (angsd -GL 1, SAMtools model) for the historical samples at the subset of 503,921 unlinked, biallelic SNPs used for PCA of contemporary genomes (see above). We then merged these samples with the contemporary Western Palearctic sample and conducted a joint PCA as described above (PCAngsd followed by `princomp`).

### Analysis of latitudinal gradient

We modeled each *pipiens* population in the Western Palearctic as a mix of northern *pipiens* and *molestus* populations using genome-wide *f3* statistics (Fig. 3A–C). Specifically, we used the `threepop` function in Treemix v1.13 (*79*) to calculate *f3*(X; *pipiens, molestus*), where X represents a focal *pipiens* population, and the *pipiens* and *molestus* reference populations came from Sweden (SWE) and Belgium (BVR), respectively. We used a block jackknife approach to obtain the standard error and compute Z-scores, dividing the genome into blocks of 500 SNPs (-k 500). A Z-score of −3 was used as a significance threshold (*79*).

We also estimated the number of derived alleles shared by focal *pipiens* populations with the same northern *pipiens* and *molestus* reference populations using Dsuite `Dtrios` (*80*), with *Cx. torrentium* as an outgroup. We specifically calculated the number of derived alleles shared with *molestus* as a fraction of those shared with either *pipiens* or *molestus*: *n*(ABBA) / (*n*(ABBA) + *n*(BBAA)) where A represents the ancestral allele and B represents the derived allele as in the tree shown in Fig. 3D.

### Distance (*Dxy*) tree inference

We inferred a distance tree for *pipiens* and *molestus* mosquitoes with >10X genome wide coverage from the full global sample based on the number of pairwise nucleotide differences (*Dxy*). Sequenced mosquitoes from early branching *Cx. pipiens* lineages native to southern Africa (named ‘juppi’ and ‘mada’), as well as *Cx. torrentium* were included as outgroups, but *Cx. quinquefasciatus* was excluded. As hybridization can confound relationships in distance trees, we used a variety of methods to identify and exclude populations or individuals that showed signs of admixture. Using *f3* tests we found that no *molestus* populations were well modeled as a mixture of *pipiens* and *molestus*, suggesting that introgression from *pipiens* into *molestus* is generally rare (fig. S3). However, a more sensitive 4-population test (Patterson’s *D*) found small yet significant signs of introgression into some Mediterranean *molestus* populations (fig. S4), which we then excluded from the tree. Identification of *pipiens* populations that have received genetic input from *molestus* is more challenging because introgression is confounded by the ancestral genetic gradient (Fig. 3). To overcome this, we used the F-branch statistics (*43*) as presented in Fig. 5A (see *Quantifying gene flow from molestus into pipiens* for details) and then excluded all *pipiens* populations that showed non-zero introgression. Finally, we excluded any *pipiens* or *molestus* individual with >2% inferred ancestry from sibling species *Cx. quinquefasciatus* based on an NGSadmix (*75*) analysis presented in the companion study (Table S1 column X). Such introgression is rare in the Western Palearctic (*81*), but extremely common in the Americas, leading to exclusion of most American samples. After filtering, we moved forward with *Dxy* tree inference for 99 *molestus*, 96 *pipiens*, and 10 individuals from outgroups.

We used pixy v.1.2.7 (*82*) to estimate pairwise genome-wide *Dxy* among the remaining samples. We included invariant accessible sites in addition to the full set of 30.6M biallelic SNPs as exclusion of invariant sites is known to generate bias (*82*). We bootstrapped genome-wide *Dxy* estimates 100 times by sampling 1Mb windows with replacement. We then built the genome-wide neighbor-joining tree as well as bootstrapped trees based on the resulting matrices of *Dxy* values using the R packages `ape` v.5.6.2 (*83*) and `ggtree` v3.6.2 (*84*).

We annotated populations in the tree based on microhabitat of origin—aboveground, belowground, or “suspected belowground”. Suspected belowground populations included one Belgian population (BVR) and one Chinese population (BEJ). The Belgian individuals were collected aboveground in a heavily industrialized zone and suspected of having escaped from a nearby tire factory. The Chinese individuals were collected trying to bite the collector inside a residential building in Tangshan, near Beijing.

### Genetic diversity (π)

We calculated genome-wide nucleotide diversity (π) for all *molestus* populations included in the *Dxy* tree analysis using pixy v.1.2.7 (*82*). A potential concern in doing so was that the eastern Mediterranean *molestus* populations, including key populations from Egypt and Israel, might have experienced introgression from *Cx. quinquefasciatus* below the 2% threshold we used for exclusion from the tree (*81*). Even a small amount of introgression from the divergent *Cx. quinquefasciatus* could inflate diversity estimates. To identify putatively introgressed genomic regions, we used Dsuite `Dtrios` to calculate *f4* admixture ratios in non-overlapping windows of fixed size (50kb, 150kb, 250kb, 500kb, 1Mb) using the following tree: (((*pipiens*, X), *quinquefasciatus*), outgroup). Reference *pipiens* and *quinquefasciatus* populations came from Sweden (SWE) and Saudi Arabia (JED). *Cx. torrentium* was used as the outgroup. The vast majority of windows in most individuals showed 0 introgression, but we observed a minor peak at *f4* ∼ 0.5 in some samples (fig. S6), likely representing the heterozygous state for introgressed haplotypes. Homozygous *quinquefasciatus* haplotypes (*f4* ∼ 1) were also sometimes present, but extremely rare. After comparing signal to noise ratios, we settled on a window size of 150kb and an *f4* cutoff of 0.2 for calling introgression (fig. S6). When computing diversity (π), we masked every 150kb locus for which *any* of the 99 *molestus* individuals showed significant introgression from *quinquefasciatus*. In total, we masked ∼5% of all 30.6M accessible sites.

### Cross-coalescent analysis of *pipiens*-*molestus* split time

To estimate the divergence time between *pipiens* and *molestus*, we carried out cross-coalescent analyses using MSMC2 following published best practices (*39*, *85*). As MSMC2 requires phased genomes, we assembled a genome phasing panel using 551 individuals with >10X coverage that represent all major geographic regions where *pipiens* and *molestus* occur (mean coverage = 19.6X, range = 10–87.3X). We considered the full set of 30.6M biallelic SNPs but further filtered out genotypes with DP < 8. We first individually phased nearby heterozygous sites based on information present in sequencing reads using HAPCUT2 (*86*). This read-based phasing alone was able to phase up to ∼90% of variants in the highest-coverage samples (range = 0.8–90.2%, median = 22.9%). We then carried out statistical phasing with the pre-phased variants across all individuals using SHAPEIT4 v2.2 (*87*) with a phase set error rate of 0.0001. To increase accuracy, we increased the MCMC iterations in SHAPEIT4 from the default value of 15 to 27 (`--mcmc-iterations 10b + 1p + 1b + 1p + 1b + 1p + 1b + 1p + 10m`), and we increased PBWT depth from the default value of 4 to 8. We phased variants on each chromosome separately.

We selected two high-coverage individuals from an Egyptian *molestus* population (ADR, 47.5X and 56X coverage) and another two from a Moroccan *pipiens* population (MAK, 56.1X and 67.5X). We first extracted phased genomes of focal individuals using BCFtools and generated chromosome-specific masks based on average coverage using bamCaller.py (*85*). We also masked every 150kb locus at which the individuals showed signs of introgression from *quinquefasciatus* (see above, *f4* > 0.2; fig. S6). We then ran MSMC2 to characterize rates of cross-coalescence within and between the two populations. The time at which the relative rate of cross-coalescence exceeded 50% was used as a point estimate of the split time (*39*). We bootstrapped MSMC2 analyses using 100 replicates of three 200 Mb ‘chromosomes’, each composed of resampled blocks of 10 Mb. To explore the robustness of our results to sample selection, we reran the analysis with an alternative Mediterranean *pipiens* population for which high-coverage genomes were available (MEG, Armenia; 50.8X and 20.7X) (fig. S7).

MSMC2 generates split time estimates in coalescent units (*85*), which can be converted to years given a taxon-specific mutation rate (*μ*) and generation time (*g*). Since *μ* and *g* have not been directly measured in natural *Cx. pipiens* populations, we used plausible, literature-based, ‘best-guess’ values, as well as biologically reasonable minima and maxima. For *μ*, we considered published data from mosquitoes and other insects and set the reasonable range at 1.0–8.0×10^−9^ (*88*–*90*). Our best-guess of *μ* was 4.85×10^−9^, taken from a recent estimate in *Ae. aegypti*, a well-studied mosquito from the same subfamily (*55*). For *g*, our best-guess was 20 days, based on a study of an autogenous *molestus* lab colony (20–21.3 days) (*91*). However, lab conditions are often better than those found in nature (e.g., unlimited food) and *pipiens* mosquitoes might be delayed in finding bloodmeals. We therefore extended the reasonable range up to 30 days. Taken together, we used the following combinations of parameters for conversion of coalescent units to our best-guess, minimum, and maximum chronological split times (Fig. 4D): *μ* = 4.85×10^−9^ and *g* = 20 (best-guess split time), *μ* = 8×10^−9^ and *g* = 20 (minimum split time), *μ* = 1×10^−9^ and *g* = 30 (maximum split time).

### Quantifying gene flow from *molestus* into *pipiens*

To quantify gene flow from *molestus* into *pipiens* across the Western Palearctic while accounting for the ancestral genetic gradient, we used Dsuite `Fbranch` to calculate branch-specific *f4* admixture ratios (*43*) (Fig. 5). We specified the tree shown in Fig. 5B and estimated gene flow into focal *pipiens* populations from the three other branches. Latitudinally varying gene flow from a northern *pipiens* population (SWE, Sweden, arrow 1) accounted for the ancestral gradient. Gene flow from a Middle Eastern *molestus* population (ADR, Egypt, arrow 2) and a northern *molestus* population (BVR, Belgium, arrow 3) allowed us to isolate genetic input from *molestus* subsequent to the split with *pipiens*.

We used a linear modeling framework to explore a potential association between *molestus* gene flow (Fig. 5D) and human population density (a proxy for urbanization). We first downloaded 30-second resolution population density data from the Gridded Population of the World v4 (*92*) and compared the effect of density on introgression when averaging density within circles of the following radii (centered around collection sites): 1, 2, 3, 4, 5, 6, 7, 8, 9, 10, 11, 12, 13, 14, 15, 16, 17, 18, 19, 20, 30, 40, 50, 60, 70, 80, 90, and 100 km. In a simple linear regression excluding three outlier populations (PAR, LND, FON, Cook’s distance > 4), human density had a significant effect using radii of 1–10 km, but not across larger distances (Fig. 5E). The model with human density averaged across a 3 km buffer explained the most variance (*R*^*2*^ = 0.21) and was used in the analysis shown in Fig. 5F. We also asked whether climate could explain additional variance in *molestus* introgression across populations by adding WorldClim2 bioclimatic variables (Bio1–19) (*93*) to the human density only model one at a time, again in a linear modeling framework. None of the bioclimatic variables significantly improved the model. Bio8 (mean temperature of the wettest quarter) was the only variable that had a marginal effect (linear model *P* = 0.09).

## Supplementary Figures

**Fig. S1.**
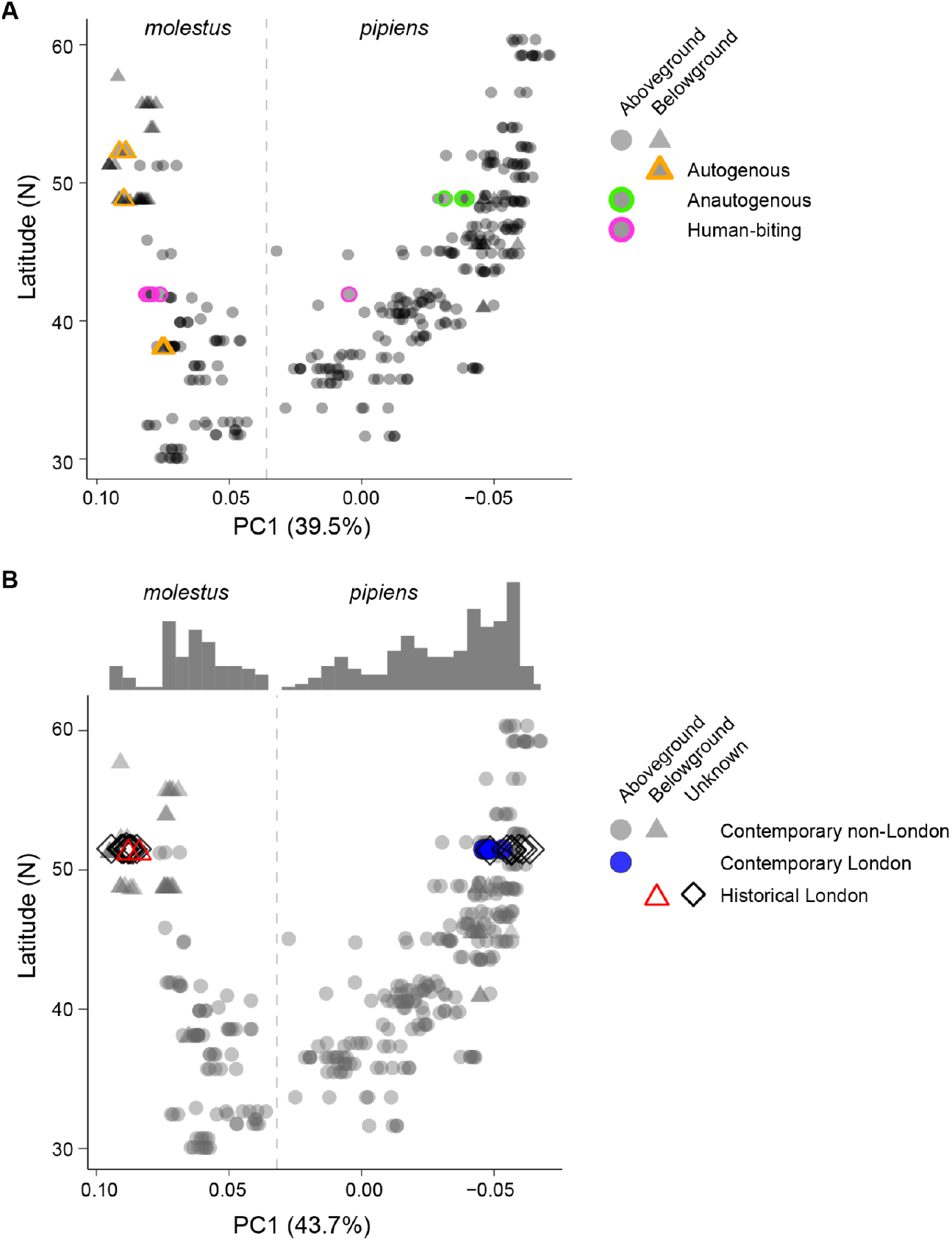
Location of phenotyped and historical samples in PCA analyses. (**A**) Individuals with known phenotypes highlighted in PC1 x latitude plot for contemporary Western Palearctic samples (same analysis as Fig. 2C; *n* = 357). The autogenous samples (able to lay eggs without a blood meal) correspond to lab strains derived from belowground sites in Amsterdam (∼52 N°) and Athens (∼38 N°), as well as field-collected samples from a flooded area on the belowground floor of a parking complex in Paris (∼49 N°). The anautogenous samples (require a blood meal for egg development) came from various outdoor locations in Paris (∼49 N°; cemetery, school, and hospital). The human-biting mosquitoes were collected trying to bite the collector in a residence in Rome (∼42 N°). (**B**) Combined analysis of genetic variation across contemporary Western Palearctic samples (*n* = 357) and historical London samples (*n* = 22, see also Fig. 2C inset). Historical samples were collected between 1940–1985 and archived in the Natural History Museum in London. See Table S1 and S2 for detailed sample metadata.

**Figure S2.**
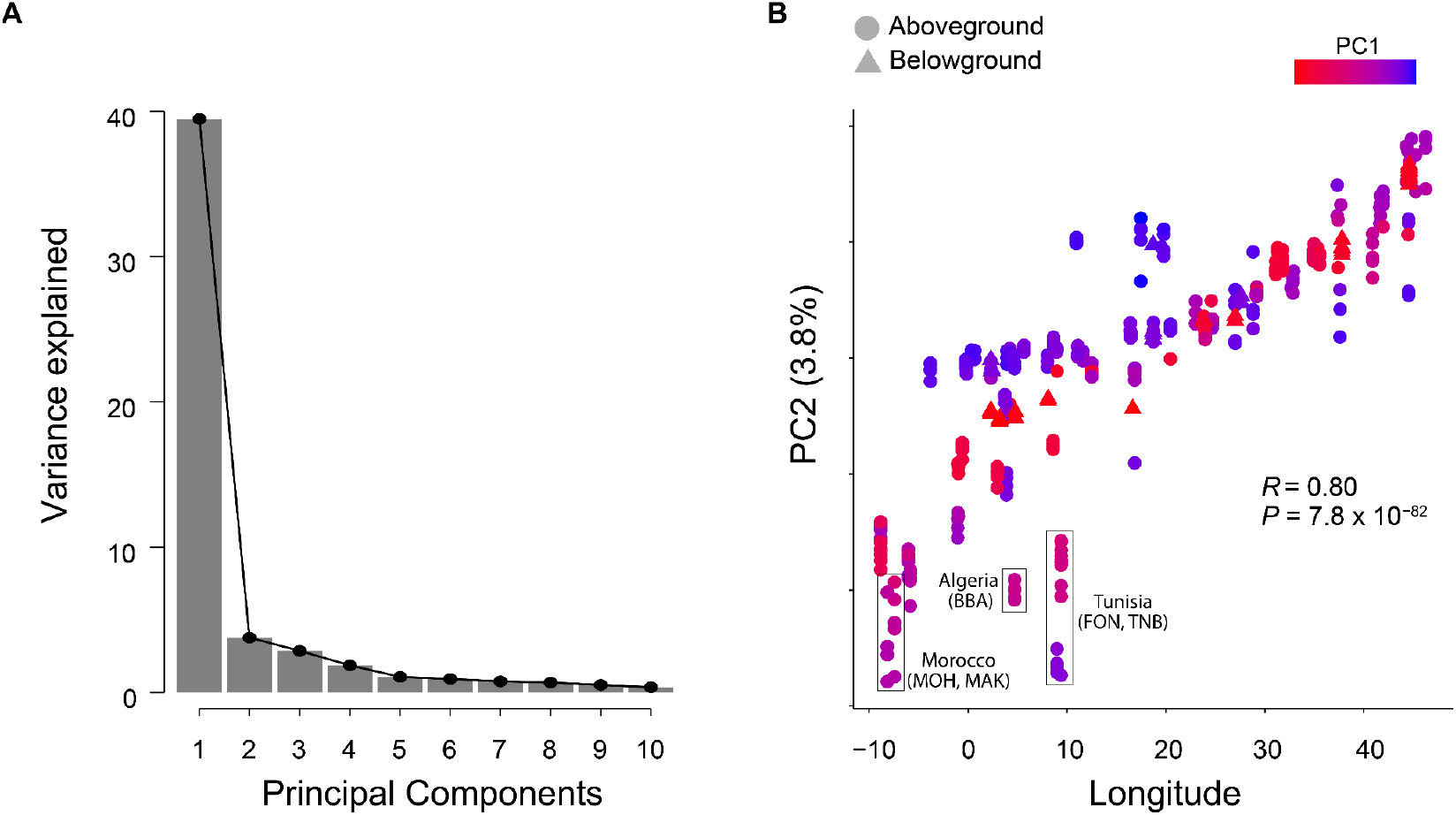
Second major axis of genetic variation across Western Palearctic samples is correlated with longitude. (**A**) Variance explained by the first 10 principal components in PCA of all Western Palearctic samples (Fig. 2B; *n* = 357 individuals). (**B**) Plot of PC2 vs Longitude showing a strong and significant correlation (Pearson’s correlation test, *R* = 0.80, *P* = 7.8 x 10^-82^). North African samples are separated from others at the same longitude, most likely due to limited gene flow across the Mediterranean Sea.

**Figure S3.**
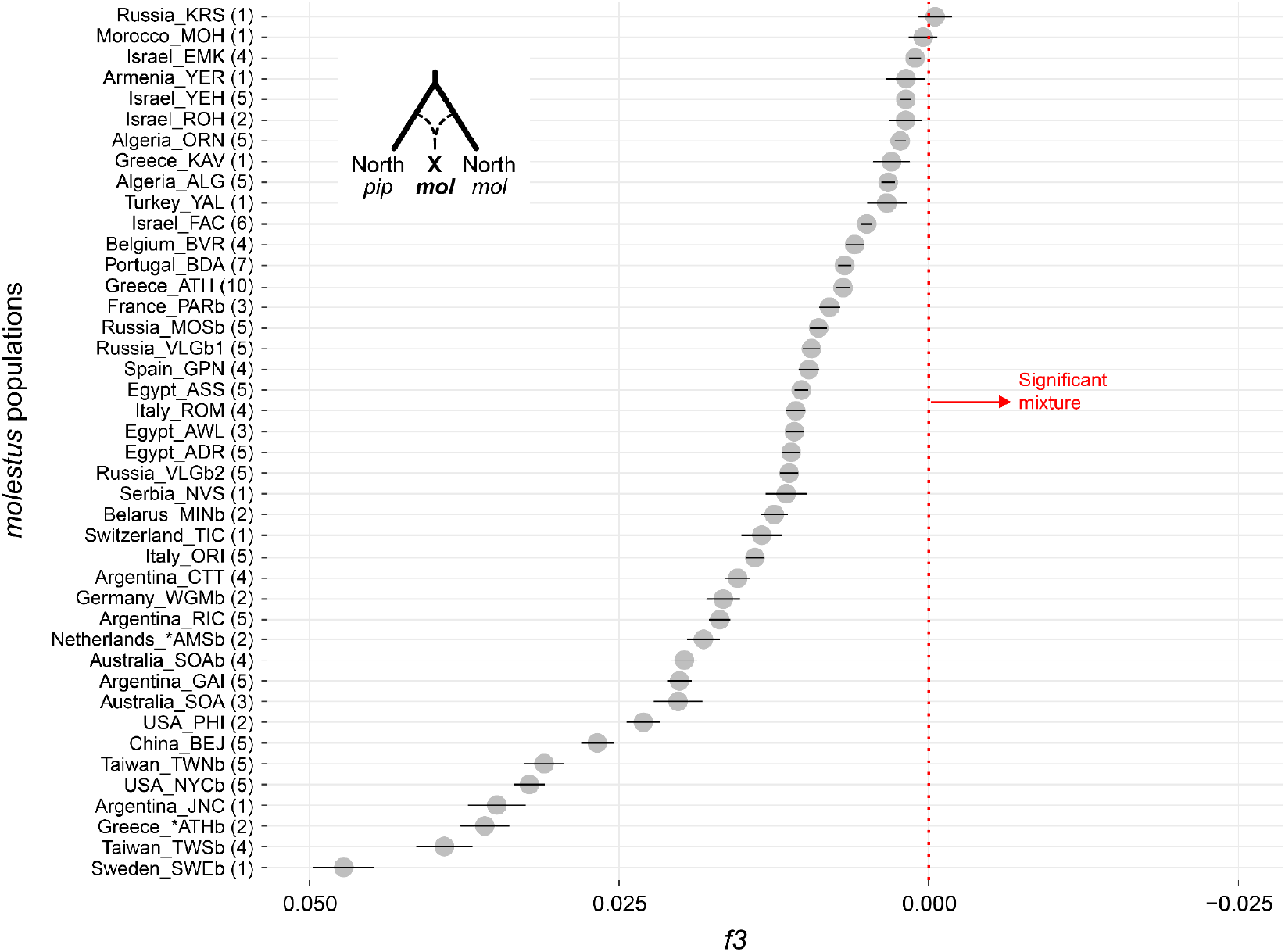
*f3* statistics for *molestus* populations modeled as a mixture of *pipiens* and *molestus* reference populations from the north. Each population was modeled as a mixture of Swedish *pipiens* (SWE) and Belgian *molestus* (BVR). Y axis labels list country, 3-letter population code (see table S1), and sample size (number of individuals). Error bars indicate 95% block-jackknife confidence intervals. None of the populations showed significant signs of mixture (*f3* < 0). This analysis includes all global *molestus* samples that showed <2% *quinquefasciatus* ancestry, inferred via NGSadmix in a companion paper (Table S1, column X). Only a few samples outside the Western Palearctic met this criterion.

**Figure S4.**
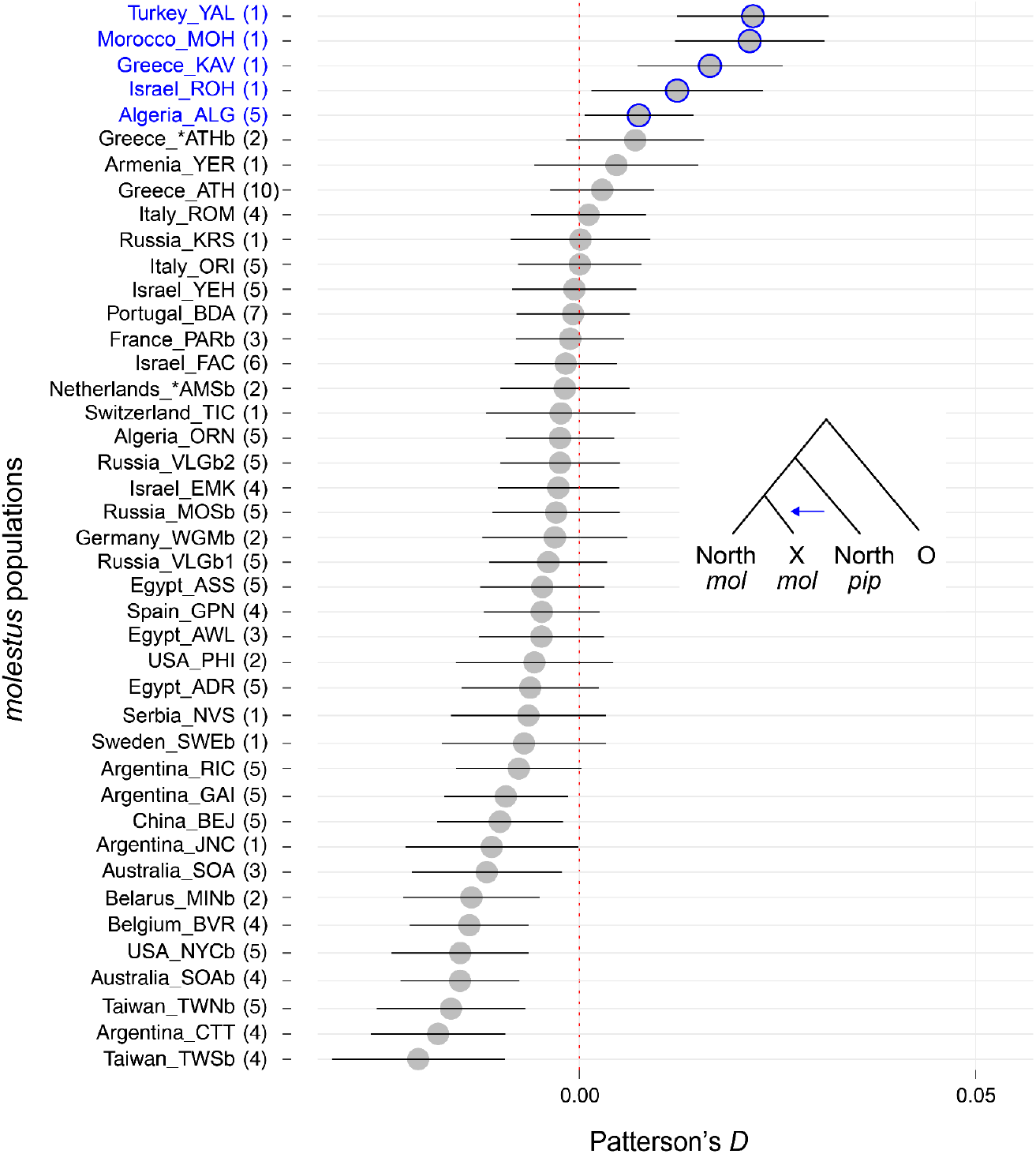
Patterson’s *D* statistics testing *molestus* populations for signs of introgression from *pipiens*. Each population was modeled as sister to Belgian *molestus* (BVR) and receiving potential genetic input from Swedish *pipiens* (SWE). Y axis labels list country, 3-letter population code (see table S1), and sample size (number of individuals). Error bars indicate 95% block-jackknife confidence intervals. Blue outlines highlight populations showing small, but significant, introgression from *pipiens* (*D*>0). These were excluded from the neighbor-joining Dxy tree in Fig. 4A and S5. This analysis includes all Western Palearctic *molestus* samples as well as any global *molestus* sample that showed <2% *quinquefasciatus* ancestry inferred via NGSadmix in a companion paper (Table S1, column X). Only a few samples outside the Western Palearctic met this criterion.

**Figure S5.**
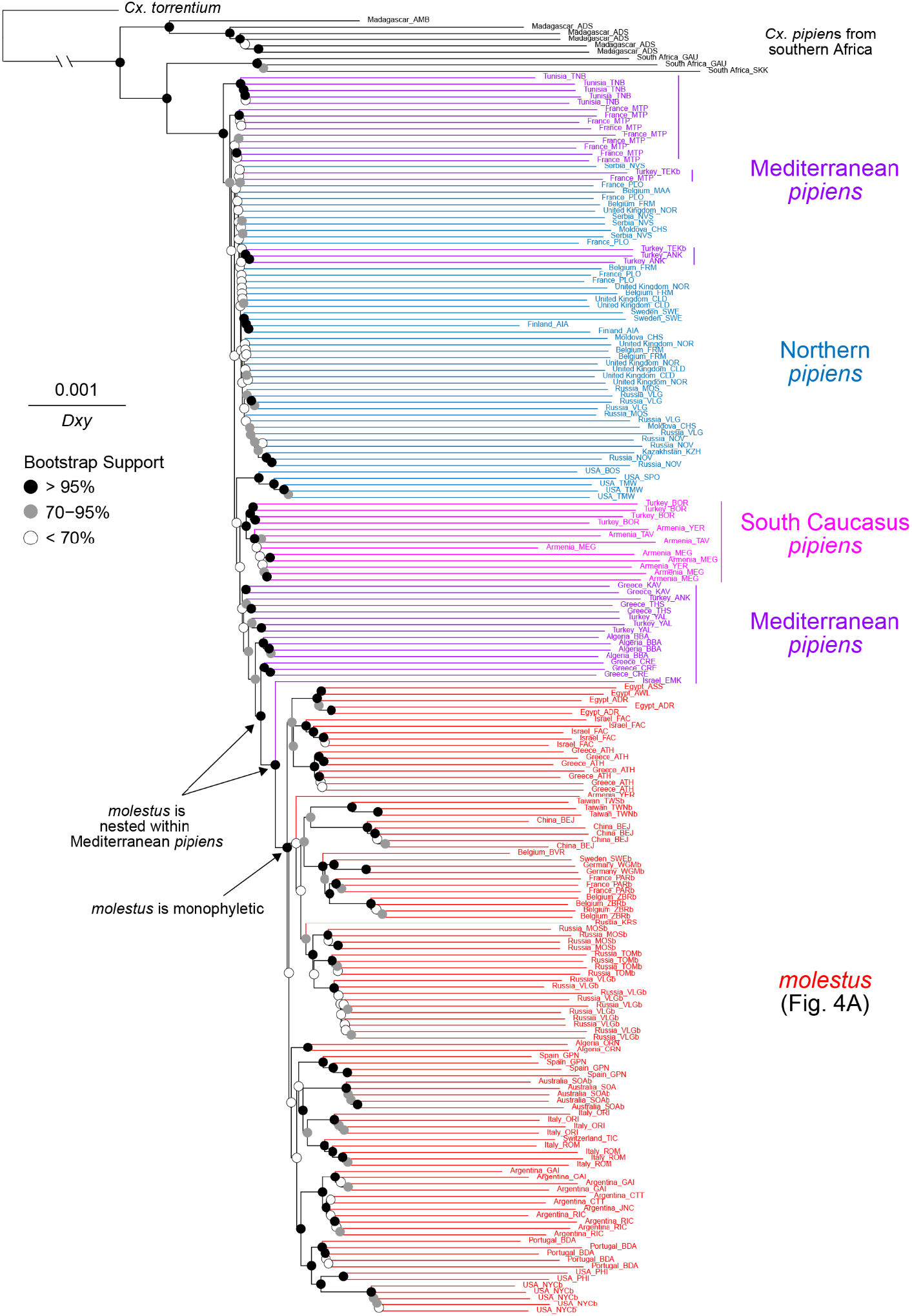
Neighbor-Joining tree based on whole genome *Dxy*, related to Figure 4A. Analysis only included individuals of ‘pure’ ancestry and >10X genome-wide coverage. *pipiens* and *molestus* individuals were deemed ‘pure’ if they showed no signs of introgression from either *Cx. quinquefasciatus* or each other (see Materials and Methods for details). A zoom version of the *molestus* clade appears in Figure 4A. *Cx. torrentium* was used as an outgroup, with branch cut short for visualization. Each tip is labeled with country of origin and population code (table S1).

**Figure S6.**
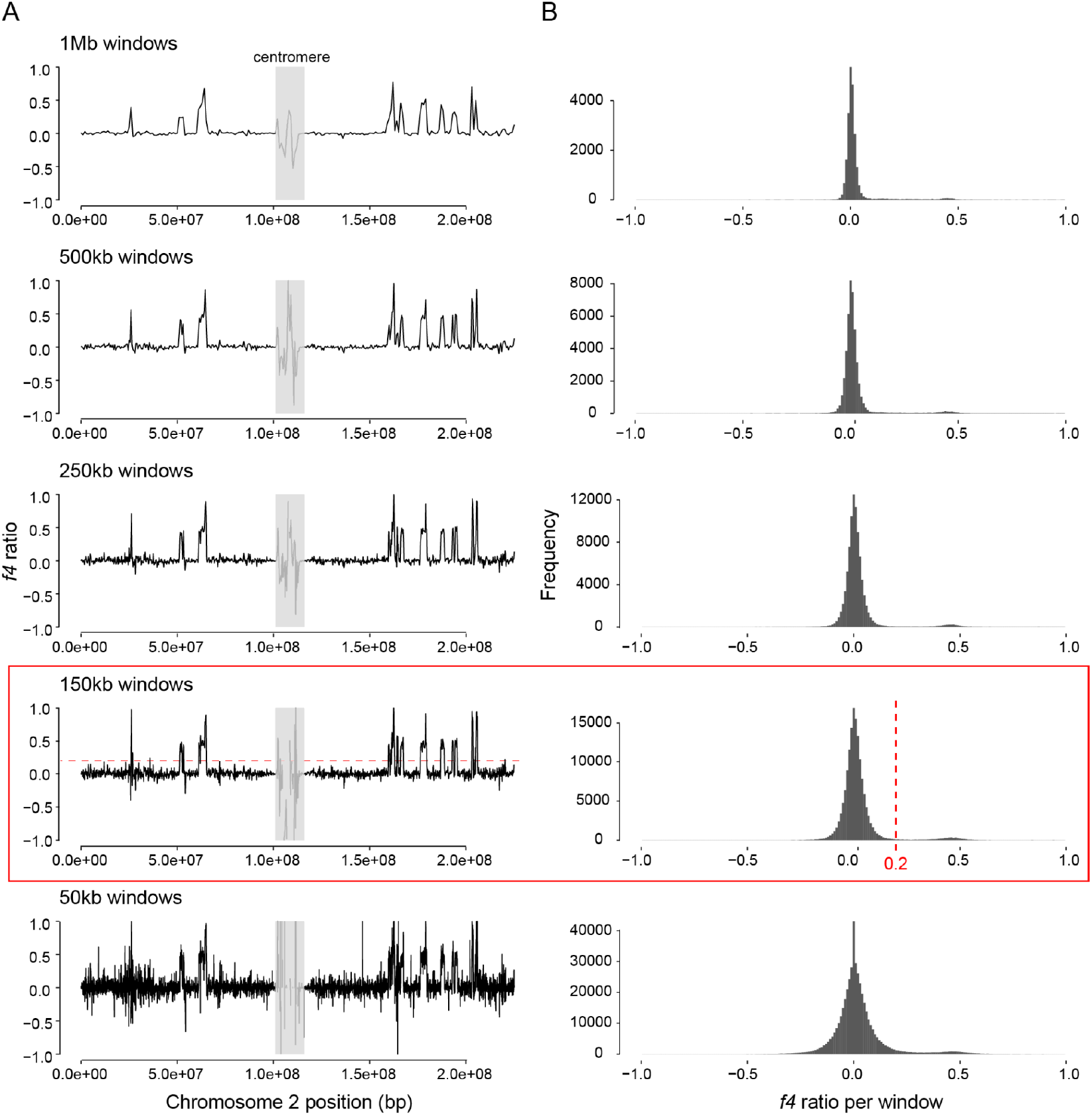
Identification of local *Cx. quinquefasciatus* introgression using *f4* ratio. We looked for stretches of introgression in the genomes of select *molestus* and *pipiens* samples so that they could be masked in analyses of genetic diversity (Fig. 4B) and cross-coalescence (Fig. 4C and S7). The figure shows example data from chromosome 2 of an Egyptian *molestus* individual (ADR1). *f4* was calculated in sliding windows of between 50kb (bottom) and 1MB (top). **A**, Full results for each window size. **B**, Summary histograms. Elevated *f4* values indicate *quinquefasciatus* introgression. Red outline and dashed lines show the chosen window size (150kb) and threshold (*f4* ratio = 0.2) chosen for masking.

**Figure S7.**
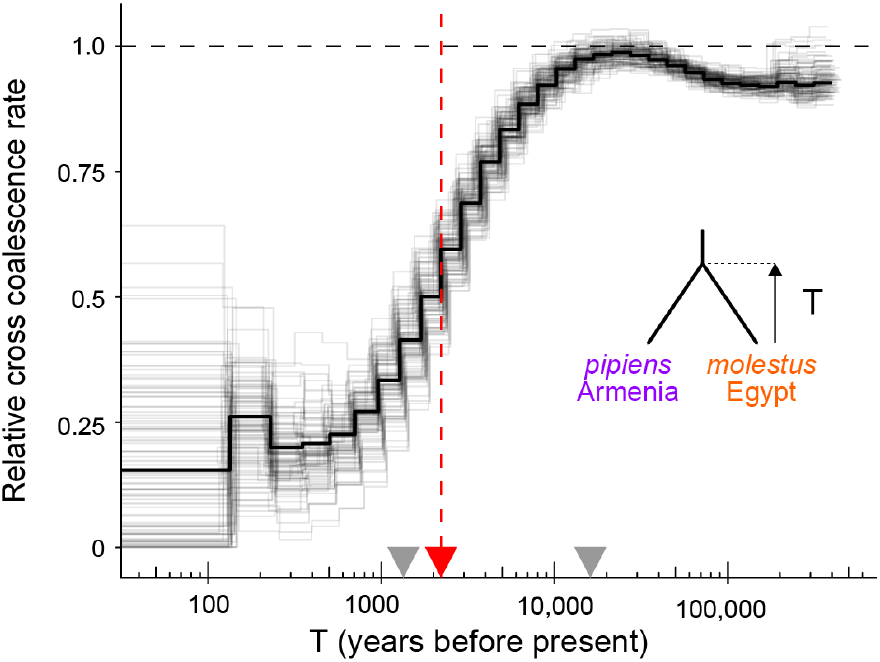
Relative cross-coalescence (rCC) rate for Armenian *pipiens* (MEG) and Egyptian *molestus* (ADR) inferred by MSMC2, related to Figure 4D. The black line indicates the genome-wide result and gray lines indicate 100 bootstrap results. Chronological time along the x-axis corresponds to a scaling factor derived from best-guess values of the *de novo* mutation rate and generation time for *Cx. pipiens (32)*. This scaling factor gives a split time (rCC = 50%) of ∼2.2K years (red dashed line and arrowhead). Minimum and maximum scaling factors (derived from biologically reasonable bounds on the mutation rate and generation time (*32*) give split times of ∼1.3K and 16.1K years (grey arrowheads).

**Figure S8.**
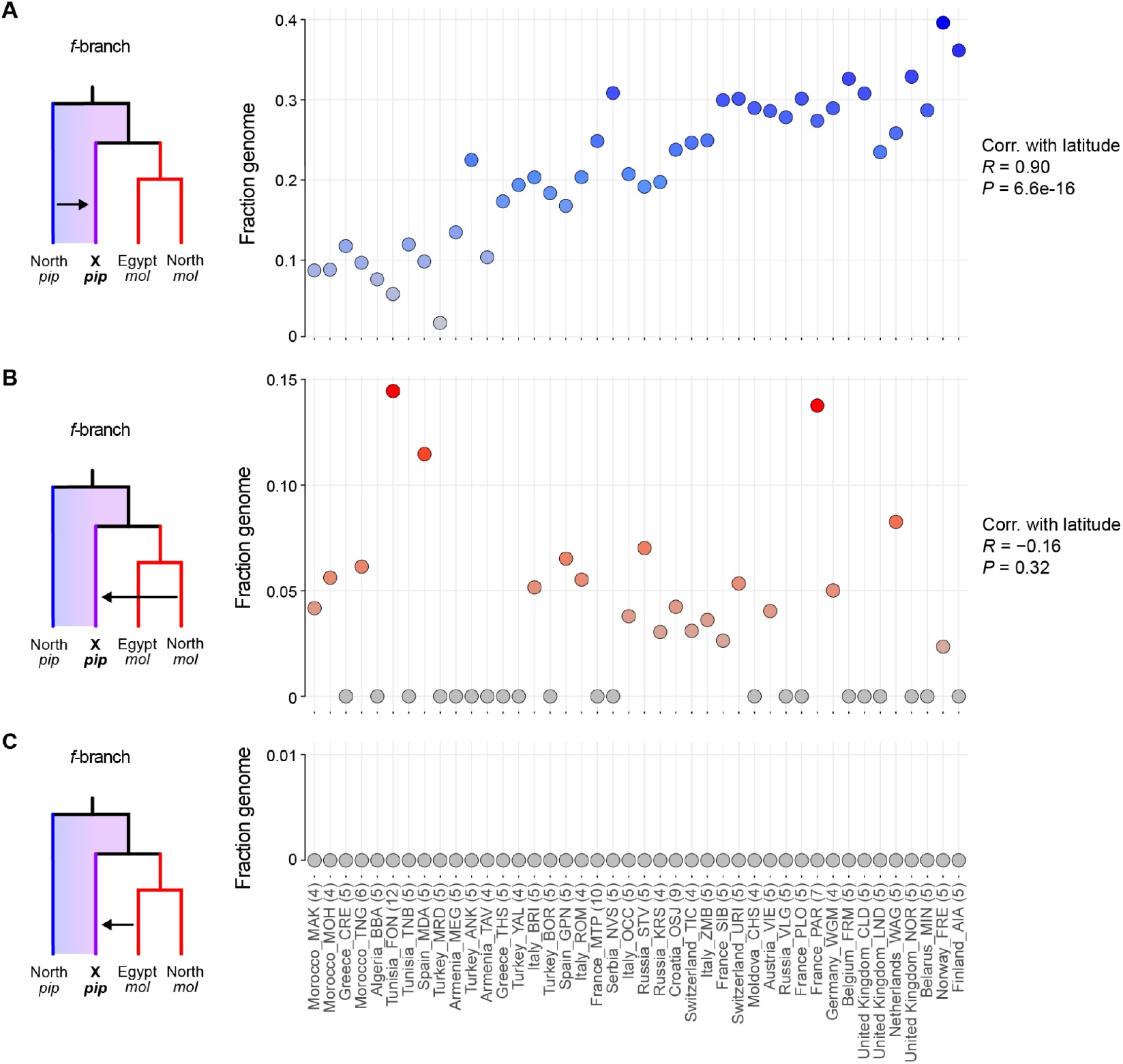
Sources of introgression into *pipiens*, related to Figure 5. Full results from *f*-branch analysis used to assess three sources of introgression into focal *pipiens* populations: (**A**) northern *pipiens* (Sweden, SWE), (**B**) northern *molestus* (Belgium, BVR), and (**C**) Egyptian *molestus* (ADR). *pipiens* populations are ordered along the *x*-axis by latitude from south (left) to north (right).

**Table S1. Sample metadata**. (Excel file)

Each row represents a mosquito sequenced for this study (*n*=790) or previously published (*n*=50, see column AA). Columns A–S contain basic collection information. Columns T and S list average sequence coverage (T) and whether or not the sample was included in the final set of 743 unrelated mosquitoes (U). Column V lists the species identity, which was usually inferred from genomic analysis but also cross-checked with information on male genitalia when available. Mosquitoes with ancestry from both *Cx. pipiens* and *Cx. quinquefasciatus* are listed as ‘hybrids’ if they showed 11–89% *Cx. quinquefasciatus* ancestry inferred using NGSadmix (column X, see below) and as one or the other ‘pure’ species otherwise. *Cx. pallens* mosquitoes from East Asia are listed as hybrids with either *Cx. quinquefasciatus* (or *Cx. pipiens*) if they showed more (or less) *Cx. quinquefasciatus* ancestry than expected given the species’ hybrid origin (cutoffs: >40% or <20% respectively). Column W lists the intraspecific *Cx. pipiens* lineage (*pipiens, molestus, juppi*, or *mada*) to which any *Cx. pipiens* or *Cx. pipiens* hybrid belongs. This inference is based on PCA and is only available for samples that passed the NGSrelate filter listed in column U. Columns X–Z show the results of an NGSadmix analysis of samples from outside sub-Saharan Africa with k=3 clusters. Note that the third cluster (Mediterranean *pipiens*/*molestus*) combines signatures of both *pipiens* and *molestus* ancestry and does not easily translate into assignment of a mosquito to the *pipiens* vs *molestus* lineage.

**Table S2. Museum specimens collected in London from 1940–1985**. (Excel file)

Each row represents one mosquito individual. Column A shows sample ID. Column B shows collection year. Column C shows locality within London, if known. Columns D and E show GPS coordinates (Latitude and Longitude). Note the coordinates are approximate, inferred from the location names on the collection note. Column F shows additional collection notes, if any. Column G shows mean genome-wide coverage after de-duplication. Column H shows inferred ecotype of each sample based on the PCA with contemporary Western Palearctic samples (fig. S1B).

